# The medaka *alg2* mutant is a model for hypo-*N*-glycosylation-associated retinitis pigmentosa

**DOI:** 10.1101/2020.08.20.260430

**Authors:** Sevinç Gücüm, Roman Sakson, Marcus Hoffmann, Valerian Grote, Lars Beedgen, Christian Thiel, Erdmann Rapp, Thomas Ruppert, Joachim Wittbrodt, Thomas Thumberger

## Abstract

Patients suffering from Congenital Disorders of Glycosylation (CDG) carry mutations in components of the evolutionarily highly conserved protein-glycosylation-machinery. Employing targeted genome editing, we modeled alleles in medaka fish based on a mutation described in an ALG2-index patient. The multisystemic phenotypes in our *alg2* model closely resembled the patient’s syndromes. Molecularly, the mutation results in a reduction of the abundance of *N*-glycans without altering the profile of glycan structures in fish as well as in patient fibroblasts. This hypo-*N*-glycosylation impacted on protein abundance in two directions. We discovered a putative compensatory upregulation of the basic glycosylation and glycoprotein processing machinery highlighting the regulatory topology of the network. Conversely, proteins of the retinal phototransduction machinery were massively downregulated in the *alg2* model. Those relate to the specific loss of rod photoreceptors that fail to be maintained in the *alg2* mutants, a condition known as retinitis pigmentosa. Transient supply of human or medaka *alg2* mRNA efficiently rescued the phenotypic spectrum and restored viability demonstrating that our *alg2* model delivers key traits for the potential treatment of the disorder.

## Introduction

Protein glycosylation is a crucial prerequisite for proper protein function during the establishment and maintenance of cells, tissues, organs and entire organisms. This post and co-translational modification is indispensable for correct protein folding and stability and thus immediately impacts on function. Strikingly, the glycosylation and function of an enormous number of target proteins is controlled by a comparably small number of proteins directing the three major routes for oligosaccharide decoration of nascent proteins, namely *N*- and *O*-linked glycosylation as well as *C*-mannosylation. Classical genetic approaches, i.e. the elimination of individual genes to study the cellular and organismal consequences, are reaching their limits since most of the mutations introduced trigger pleiotropic consequences and are (embryonically) lethal. When glycosylation is affected in patients, those suffer from Congenital Disorders of Glycosylation (CDG) and develop severe clinical multisystemic phenotypes. Improper glycosylation affects, for instance, neuronal development as reflected by defects in myelination and brain formation (Hennet & Cabalzar, 2015) as well as muscle function (Muntoni *et al*, 2008). As part of the brain, the eye is affected in its development and function, as apparent by developing strabismus, cataract, coloboma formation and the (progressive) neurodegenerative disease retinitis pigmentosa (Cantagrel *et al*, 2010; Morava *et al*, 2010).

For *N*-linked glycosylation, the initial oligosaccharides are assembled at the membrane of the endoplasmic reticulum (ER) with the early steps taking place on the cytosolic face. During this lipid-linked oligosaccharide (LLO) synthesis, sequential addition of two *N*-acetylglycosamines (GlcNAc), nine mannoses (Man) and three glucoses (Glc) to the LLO-precursor is catalyzed by discrete Asparagine-Linked Glycosylation (ALG) enzymes. The Alpha-1,3/1,6-mannosyltransferase (ALG2) however has an unusual dual feature, as it creates the first branching *N*-glycan structure by adding mannoses to positions 3 and 6 of the Man1GlcNAc2-PP-Dol (Li *et al*, 2018; Thiel *et al*, 2003). Following the addition of two more mannoses, the LLO-precursor is internalized via a flippase mechanism onto the inner leaflet of the ER and more monosaccharides are added by several other ALG enzymes employing dolichol-linked (Dol) sugar donors. Finally, an oligosaccharyltransferase catalyzes the transfer of the mature Glc3Man9GlcNAc2-PP-Dol from the lipid to an asparagine acceptor amino acid of the NXS/T (X not Proline) motif on the nascent polypeptide (Breitling & Aebi, 2013). After passing the quality control in the ER, correctly folded glycoproteins eventually exit the ER to the Golgi, where the *N*-linked glycan may undergo further modifications to form complex glycan structures (Helenius & Aebi, 2004). The ALG enzymes and the resulting glycan structures are evolutionarily highly conserved across almost all eukaryotes, underpinning their crucial importance (Park & Zhang, 2011).

Since severe mutations in the corresponding genes are lethal, the majority of CDG patients suffer from hypomorphic mutations that result in reduced protein functionality. ALG2 patients (CDG type Ii) appear normal at birth, but they subsequently exhibit prominent deficits in muscles, eye and brain. In particular iris coloboma, neuronal symptoms such as hypomyelination and intractable seizures as well as mental retardation have been described (Cossins *et al*, 2013; Monies *et al*, 2016; Thiel *et al*, 2003). In addition patients suffer from hepatomegaly and coagulopathy (Haeuptle & Hennet, 2009; Varki *et al*, 2019). ALG2 is highly sensitive to alterations and its full functionality at the basis of the *N*-glycosylation cascade is crucial. Consequently, most mutations result in its loss-of-function, explaining the low number of ALG2 patients (to date 9 patients from 5 families are reported) (Cossins *et al*, 2013; Monies *et al*, 2016; Thiel *et al*, 2003). To study the function of glycosyltransferases and the consequences of their mutation, predominantly mouse models have been employed. The absence of *N*-linked glycosylation results in early embryonic, pre-implantation lethality (Matthijs *et al*, 1998; Med2002; Thiel *et al*, 2006) and (early onset) phenotypes can only be observed upon sacrificing the heterozygous mothers. Interestingly, a mouse model based on a hypomorphic mutation in the *mannose phosphate isomerase* (*MPI)* detected in patients is viable (Sharma *et al*, 2014), arguing for establishing mutant alleles in model systems on the molecular details of CDG patients.

The extrauterine development of teleost models (zebrafish, *Danio rerio*; medaka, *Oryzias latipes*) facilitates addressing the developmental consequences of reduced glycosylation in the organismal context *in vivo*. Upon precise genome editing employing the CRISPR/Cas9 toolbox (Gutierrez-Triana *et al*, 2018; Stemmer *et al*, 2015) mutations can be modelled based on the well-described human alleles. We addressed the initial steps of the *N*-glycosylation pathway and generated viable hypomorphs of *alg2*. Employing CRISPR/Cas9-mediated genome editing in medaka, we introduced mutations at a site orthologous to the site of mutation described in a human ALG2 index patient (Thiel *et al*, 2003). Our small animal model resembled the human ALG2-CDG phenotype. We validated the hypomorphic nature of the alleles generated. Proteomic analysis uncovered compensatory routes to maintain minimal Alg2 activity as well as protein targets that provide a molecular basis for the observed complex phenotype. This is in particular true for the eye where several proteins implicated in retinitis pigmentosa were severely misregulated. Strikingly, a progressive elimination of rod cells in the photoreceptor layer of the retina was observed in the Alg2 fish model as predicted by the Alg2 targets identified. The mutant phenotype is efficiently rescued by additional Alg2 activity provided during embryonic development.

## Results

### Medaka *alg2* variants recapitulated the symptoms of CDG type I

Since the complete loss-of-function of glycosylation related proteins is lethal, we took advantage of the described hypomorphic mutation in a human ALG2 index patient (c.1040delG that translates to p.G347Vfs*26; Fig 1A and A’; (Thiel *et al*, 2003)) to introduce a mutation at the corresponding position of the orthologous medaka *alg2* gene. Employing targeted CRISPR/Cas9 genome editing (Fig 1A and Fig EV1), we used a single-stranded oligodeoxynucleotide (ssODN) integration as well as indel formation. Survival of the genome-edited embryos was used to select for viable Alg2 variants in medaka. We established three different alleles in stable fish lines (predicted resulting proteins depicted in Fig 1A’; genomic sequences see Fig EV1) that show alterations corresponding to the site of mutation in the ALG2 index patient (dashed line, Fig 1A; (Thiel *et al*, 2003)). The ssODN edited allele displayed an early premature STOP codon (XM_004077863:c.1006-1011delinsTAAGG that translates to XP_004077911:p.G336*). A second allele represented an in-frame deletion, lacking six nucleotides at the C-terminus (XM_004077863:c.999-1004del that translates to XP_004077911:p.N334_S335del). A third allele harbored a large deletion at the C-terminus (XM_004077863:c.1002_1218+5delinsTCTG that translates to XP_004077911:p.(S335Lfs*8)). All three alleles displayed fully comparable phenotypes under homozygosity, yet for the rest of the study, we focused on the precisely edited p.G336* variant.

Mutant analysis did not reveal any apparent change in gross morphology prior to stage 39 (Iwamatsu, 2004). Similar to the multisystemic effects reported for human ALG2 patients, we observed multisystemic phenotypes with late onset during embryonic development that progressed quickly (Fig 1B). The first recognizable phenotypes of homozygous mutant embryos were reduced blood flow, followed by blood clogging and eventually a complete arrest of blood circulation. The initially well-formed two chambered heart became secondarily tubular, vessels thinned out and edemas formed around the heart and the eyes. The mutant embryos failed to engulf the yolk and a prominent yolk sac was still apparent after hatching. The air-inflated swim bladder seen in wild type (wt) hatchlings (Fig 1C) was not detectable in the mutants. In contrast to the overall comparable size of the head, trunk and tail, the eyes were slightly smaller and craniofacial dysmorphisms resulted in a severely shortened snout (Fig 1B and 1D, black bracket). The reduction in jaw size was caused by significantly shorter Meckel’s cartilage (p = 0.02), palatoquadrate (p = 0.03) and ceratohyal (p < 0.001) cartilages compared to stage-matched wt siblings (Fig1 D to F). The multisystemic phenotypes progressed quickly and resulted in lethality of homozygous mutants at 2-3 days post hatching (dph). Interestingly, the interindividual phenotypic variance of the homozygous hatchlings was minute.

**Figure 1.**
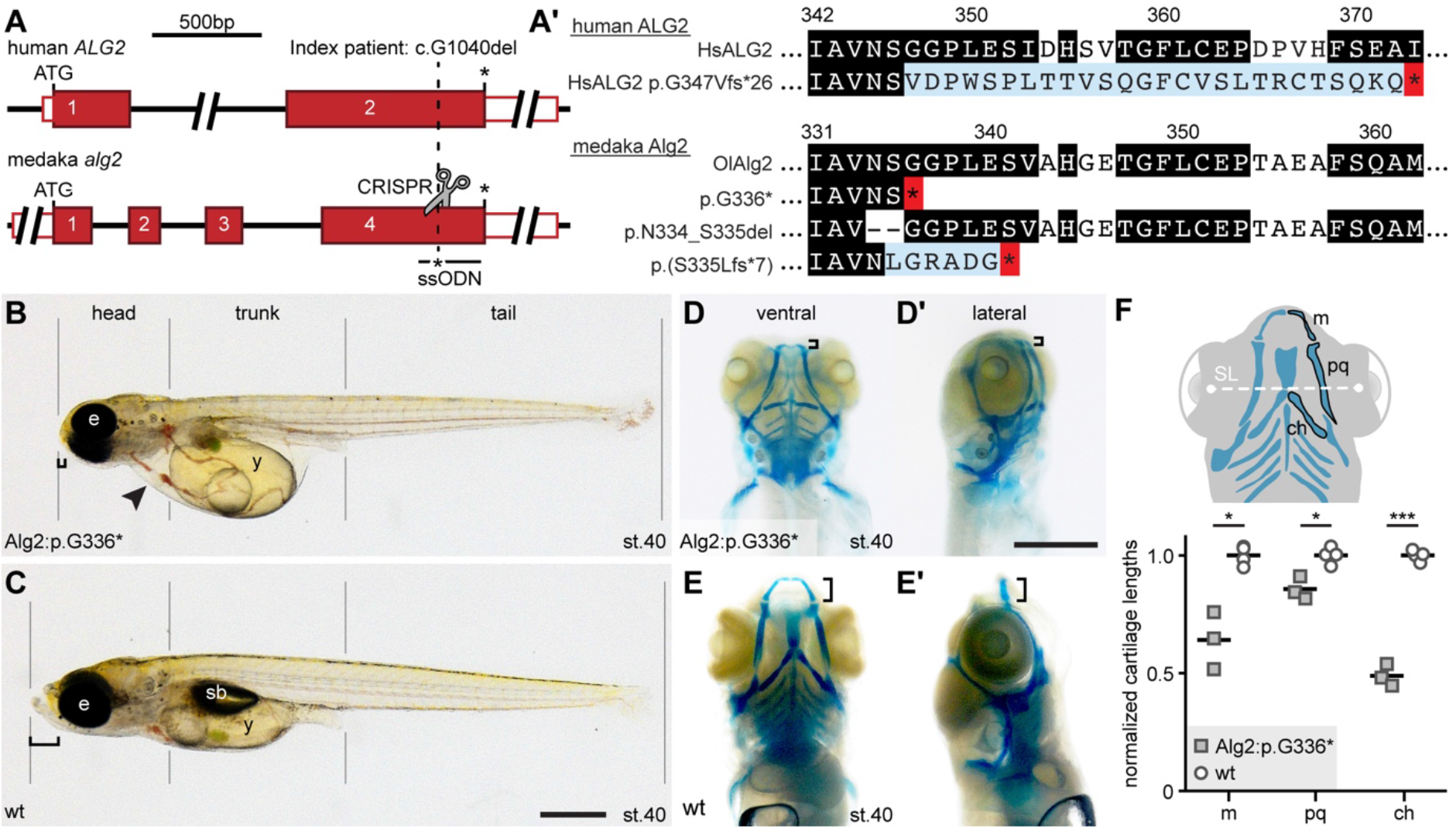
Homozygous medaka *alg2* mutants recapitulate ALG2-CDG multisystemic phenotype. A) Schematic representation of human *ALG2* and the medaka orthologous *alg2* locus. Site of ALG2 index patient mutation (c.1040delG) and targeted CRISPR genome editing (scissors) in medaka indicated (dashed line). Single-stranded oligodeoxynucleotide (ssODN) donor for introduction of premature STOP (*) given. Red box, coding exon; white box, untranslated region. A’) Predicted protein variants at the given position of human ALG2 index patient and corresponding sites of the stable mutant fish lines. Black, highly conserved amino acids (AA); blue, different AA resulting from underlying frame shift; red, premature STOP. Nomenclature of protein variants according to (Dunnen *et al*, 2016). For genomic sequences of medaka *alg2* mutant alleles see Figure EV1. B, C) Representative phenotype of Alg2:p.G336* mutant (B) and corresponding wild type (wt, C) at stage 40. Note in mutant: craniofacial defects including prominently shortened snout (bracket), persistent yolk sac (y), non-inflated swim bladder (sb), secondary tubular heart (arrow) and clogging of blood, slightly smaller eyes (e). D-E’) Alcian blue staining of Alg2:p.G336* mutant (D-D’) and wt (E-E’) embryos at stage 40 reveal dramatic reduction of the craniofacial cartilages (brackets). ventral view in D, E; lateral view D’, E’. F) Schematic representation (upper panel, ventral view) of craniofacial cartilages and comparison (lower panel) of measured cartilage lengths (m, Meckel’s cartilage; pq, palatoquadrate; ch, ceratohyal) normalized to distance between lenses (standard length, SL) and mean of wt reveals significant reduction of all three cartilage structures in mutant compared to wt (two-tailed nonparametric Student’s T-test; * < 0.05; *** < 0.001). Scale bars: 0.5 mm

Overall, the medaka model presented here successfully recapitulated the symptoms described for type I CDG. All three established mutant lines that harbored mutations at the site corresponding to the mutation in the ALG2 index patient resulted in identical phenotypes. Since the CDG patients suffer from hypoglycosylation (Thiel *et al*, 2003), we investigated whether this also holds true for the medaka mutant.

### Medaka Alg2:p.G336* variant resulted in hypo-*N*-glycosylation

To address the overall *N*-glycan occupancy of proteins, we employed the specific glycan-binding capacity of the lectins Concanavalin A (Con A) to detect mannose and wheat germ agglutinin (WGA) to detect *N*-acetylglucosamine. Lectin blots on proteins extracted from *alg2:p.G336\** mutant medaka hatchlings showed reduced levels of mannose (72%, ConA) and *N*-acetylglucosamine (83%; WGA) compared to the wt siblings (Fig 2A). The analysis of fibroblasts derived from the ALG2 index patient by the same approach did not show this tendency as the levels of mannose (102 %; Con A) and *N*-acetylglucosamine (94 %; WGA blot) were comparable (Fig 2B).

**Figure 2.**
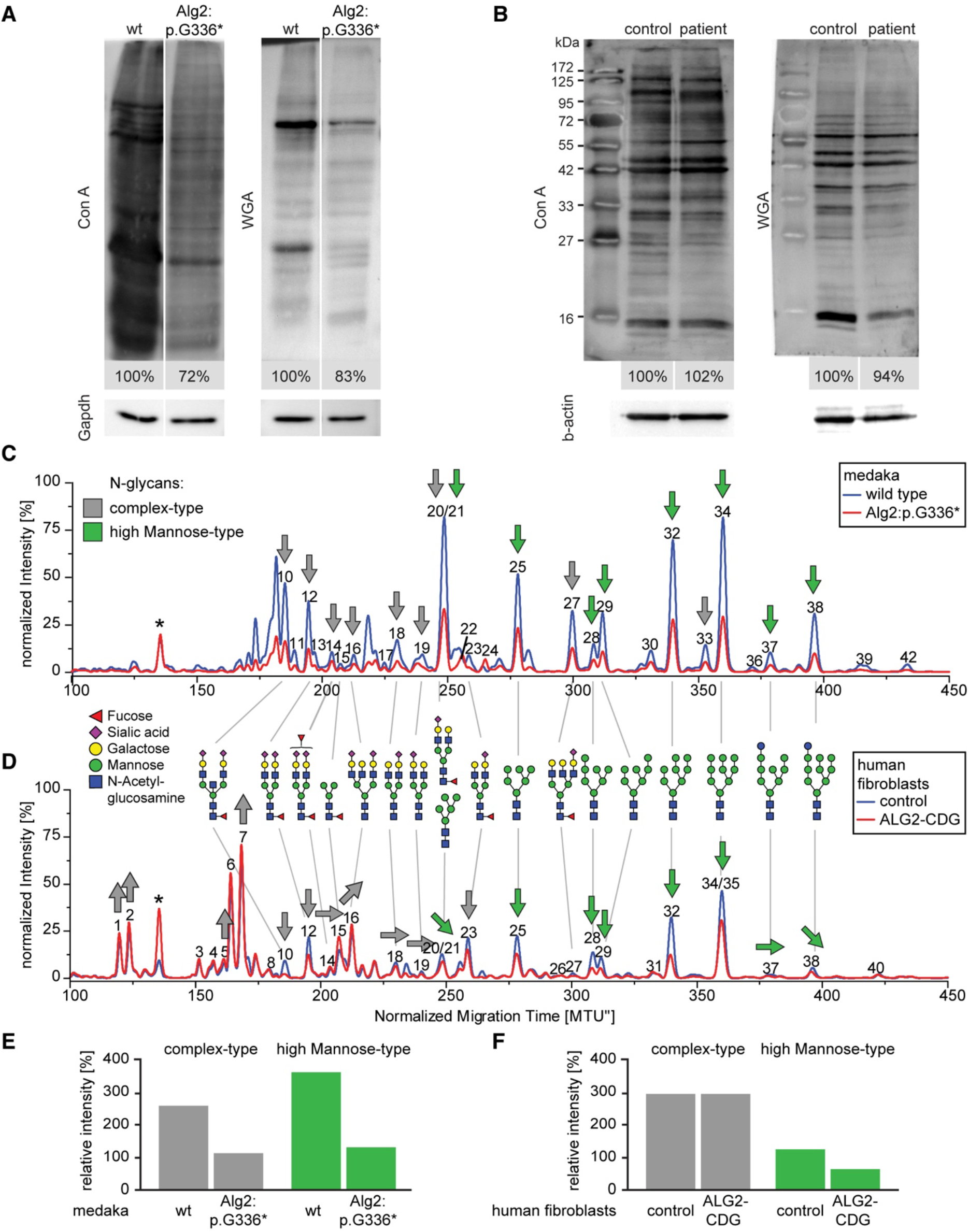
Medaka Alg2:p.G336* and human ALG2:p.G347Vfs*26 are hypomorphic variants. A, B) Lectin blots on total protein lysates of medaka wild type (wt) and Alg2:p.G336* mutants (A) as well as human control and CDG-patient fibroblasts (B). Intensity values normalized to internal loading control (Gapdh / β-actin) and wild type or control samples. Concanavalin A, Con A; Wheat Germ Agglutinin, WGA. C, D) xCGE-LIF-generated *N*-glycan fingerprints comparing medaka wt (blue) to Alg2:p.G336* mutants (red; C) and human fibroblast control (blue) to ALG2 index patient fibroblast (red; D). Asterisk indicates standard for migration time (MTU) normalization. Trends in observed quantitative changes for individual *N*-glycan structures are indicated by grey (complex-type) and green (high Mannose-type) arrows, direction indicating up or downregulation. Peak numbers refer to identified *N*-glycan structures (see list of all identified *N*-glycans Table EV1). E, F) Quantification of complex type (grey) and high Mannose-type (green) *N*-glycan structures in medaka wt versus mutant (E) and control versus patient fibroblast (F).

To further address how the medaka Alg2:p.G336* variant and fibroblasts of the ALG2 index patient impacted on the *N*-glycan occupancy, we performed an extended *N*-glycan analysis by multiplexed capillary gel electrophoresis with laser-induced fluorescence detection (xCGE-LIF). This approach enables the identification and relative quantification of individual *N*-glycan structures via normalization to an internal standard (Fig EV2), allowing the direct quantitative comparison between samples of different origin like fish and human (Hennig *et al*, 2016). The evolutionary conservation of the complex glycosylation machinery resulted in a high structural similarity of the *N*-glycan profile of wt medaka hatchlings and control human fibroblasts. In the medaka wt *N*-glycan fingerprint (Fig 2C, blue), 57 % represented complex-type *N*-glycans that were predominantly Neu5Ac-sialylated. The remaining 41 % of the *N*-glycome comprised high-mannose-type *N*-glycans ranging from Man5 to Man9 structures and hybrid-type *N*-glycans were present with only 2.5 %. In contrast, in the Alg2:p.G366* *N*-glycan fingerprint (Fig 2C, red) a severe hypo-*N*-glycosylation was apparent by an overall reduction of 61 % in the levels of *N*-glycosylation (Fig 2E). This hypoglycosylation affected all *N*-glycan structures to a comparable extent with only minor qualitative differences.

The *N*-glycan fingerprint of the human fibroblast control sample (Fig 2D, blue) revealed that, similar to fish, 68 % of the *N*-glycome was composed of complex type *N*-glycans with a substantial proportion of multi-antennary structures. The remainder consisted of high-mannose type *N*-glycans (29 %) and hybrid-type structures (3 %). Similar to the medaka samples, the majority of complex-type *N*-glycans in the human fibroblasts were characterized by a terminal sialylation. The *N*-glycan fingerprint from fibroblasts of the human ALG2 index patient (Fig 2D, red) showed few qualitative changes in its *N*-glycome compared to the control. Consistent with our findings in the medaka animal model, the human fibroblasts of the ALG2 index patient revealed a general hypo-*N*-glycosylation, although less pronounced. High-mannose-type glycosylation was reduced by 48 %, while complex type *N*-glycans showed only marginal changes (0.5 %; Fig 2F).

Besides hypo-*N*-glycosylation, overall sialylation and fucosylation levels were reduced in the medaka animal model as well as in the fibroblasts of the human patient. Our analysis revealed that both, the human *ALG2:p.G347Vfs*26* and the medaka *alg2:p.G336\** alleles resulted in a severe hypo-*N*-glycosylation and we accordingly use *alg2^hypo^* as a proxy for the medaka homozygous mutant.

Glycosylation is known to affect protein stability, we therefore investigated the consequences of the hypo-*N*-glycosylation in *alg2^hypo^* on the proteome composition.

### Hypo-*N*-glycosylation resulted in upregulation of proteins involved in glycosylation and downregulation of the phototransduction pathway

To address potential changes in the proteome, we performed an unbiased proteomics analysis. We analyzed total protein extracts of *alg2^hypo^* and wt hatchlings (stage 40, de-yolked, three replicates with six hatchlings each) upon isotopic labelling on the peptide level by dimethylation (Boersema *et al*, 2009). Interestingly, the differential proteome analysis did reveal very distinct differences, but not a global regulation of protein levels. Analysis of the proteomes of *alg2^hypo^* mutant fish (Fig 3A-C) uncovered 15 proteins with a significant (p < 0.05) differential abundance differing more than 2 fold (Fig 3B) in comparison to wt controls.

**Figure 3.**
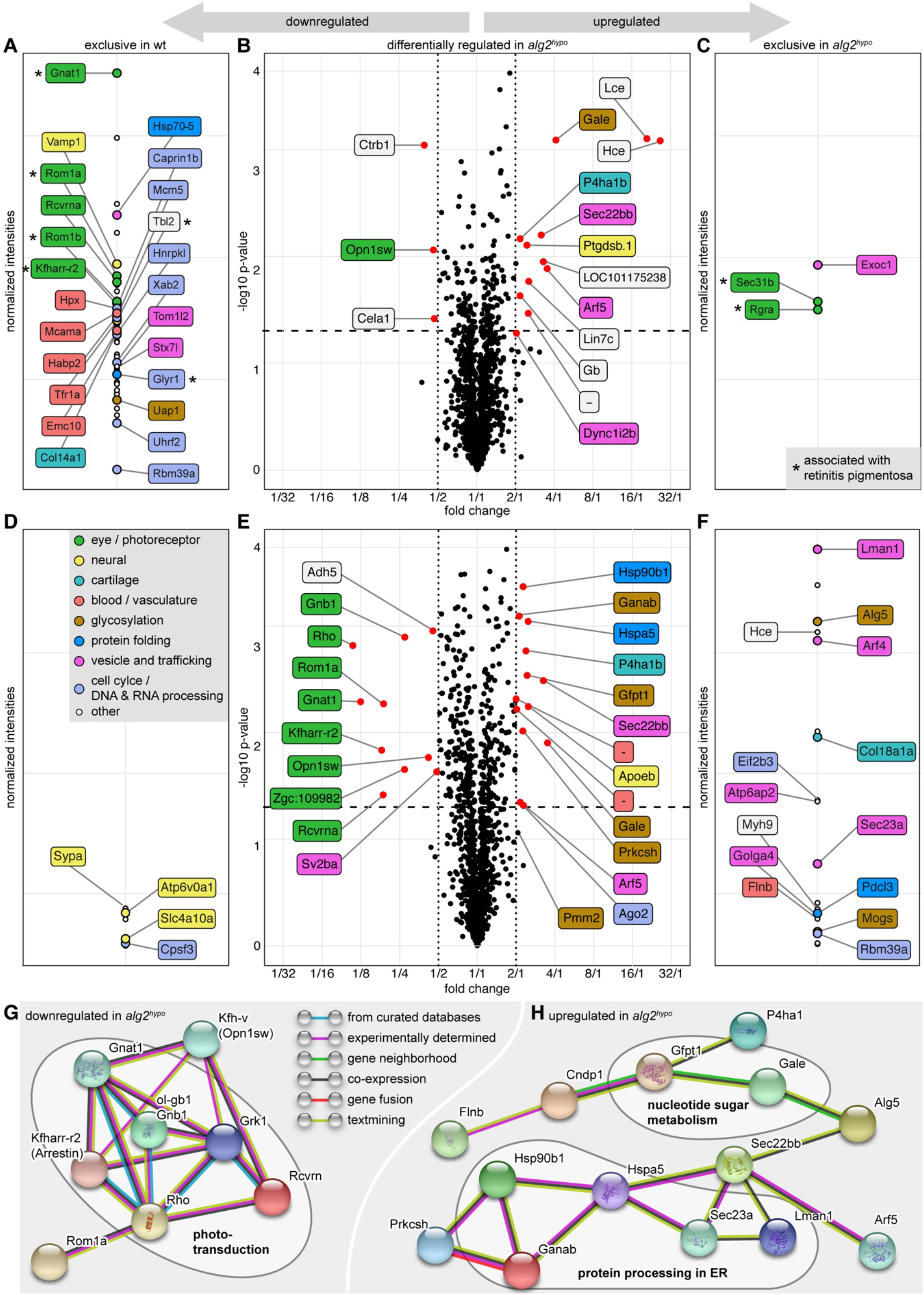
*alg2^hypo^* mutants show strong reduction of proteins involved in phototransduction and upregulation of glycosylation machinery. Unbiased mass spectrometry of chemically labelled peptides derived from total protein lysates of deyolked wild type (wt) and *alg2^hypo^* whole hatchlings (A-C) and enucleated eyes (D-H) at stage 40. Coloured labelling and grouping according to (predicted) protein function retrieved by text-mining. A, D) Intensity plots of proteins exclusively detected in wt samples amounted to 56 proteins from whole hatchling samples (A) and 7 from enucleated eye samples (D). B, E) Volcano plot of protein abundance comparison between wt and *alg2^hypo^* reveals no major regulation in protein abundance, yet distinct changes. Proteins with significant (p < 0.05, dashed line) and more than 2 fold change (dotted lines) indicated (red dots) and labelled. C, F) Intensity plot of proteins exclusively detected in *alg2^hypo^* mutant samples amounted to 3 proteins from whole hatchling samples (C), 19 proteins from enucleated eye samples (F) exclusively detected in *alg2^hypo^* samples. G, H) Network analysis of functional associations and/or interactions of all regulated and exclusive proteins in eye samples. Different clusters of regulated proteins were identified using the STRING software (medium confidence score, disconnected nodes hidden; (Szklarczyk *et al*, 2019)). Among the downregulated proteins, photoreceptor-related players of the phototransduction pathway were overrepresented (G). Among the upregulated ones, proteins involved in protein processing in the ER and of the nucleotide sugar metabolism pathways were pronounced (H). The color lines linking different proteins represent the types of evidence (blue: from curated databases; purple: experimentally determined; green: gene neighborhood; black: co-expression; red: gene fusion; light green: text mining). For full proteomics data see Table EV2. Note: asterisks in A and C mark proteins associated with retinitis pigmentosa.

Among the twelve proteins upregulated in the *alg2^hypo^* mutant fish (Fig 3B) we identified the secreted metalloendopeptidases low (lce) and high (hce) choriolytic enzymes (19 and 22 fold upregulation, respectively) as the top hits followed by UDP-glucose 4-epimerase (Gale). The latter catalyzes two distinct interconverting reactions: UDP-galactose (UDP-Gal) from and to UDP-Glc as well as *N*-acetylgalactosamine (GalNAc) from and to GlcNAc (Daenzer *et al*, 2012), all four nucleotide sugars serve as substrates for glycan synthesis. The remaining upregulated proteins are related to endopeptidases or play a role in vesicle transport, heme binding, or collagen synthesis. The three downregulated proteins were a putative violet sensitive opsin and two proteins containing peptidase S1 domain (Fig 3B). In addition, a larger number of proteins were either present in wt or mutant samples only (Table EV2). These exclusive hits amounted to 56 in wt (Fig 3A) and three in the *alg2^hypo^* samples (Fig 3C). Those proteins group into classes involved in vesicle trafficking, regulation of blood and circulatory system, protein folding as well as proteins with described and predicted functionality in neuronal and especially eye and photoreceptor related activity. Strikingly, eight of these 59 candidates are associated with retinitis pigmentosa (human orthologs in parentheses): Gnat1 (GNAT1), Rom1a and Rom1b (ROM1), Kfharr-R2 (SAG, ARRESTIN, RP47), Rgra (RGR, RP44), Tbl2 (TBL2), Sec31b (SEC31B) and Glyr1 (GLYR1) (Ba-Abbad *et al*, 2018; Carrigan *et al*, 2016; Conley *et al*, 2017; Sullivan *et al*, 2017).

To further refine the relative abundance of eye-related proteins, we performed an unbiased proteomics approach on enucleated eyes (four replicates, 30 eyes each) of *alg2^hypo^* and wt hatchlings at stage 40. This analysis identified 23 differentially expressed proteins (p < 0.05) with more than 2 fold change and 26 proteins exclusively found in wt or mutant samples (Fig 3D to 3F), while global protein levels were not changed. A protein-protein interaction analysis (STRING software v11.0; (Szklarczyk *et al*, 2019)) on the regulated and exclusively found proteins revealed distinct networks affected by hypo-*N*-glycosylation. For the proteins that were downregulated in the *alg2^hypo^* mutant, players of the phototransduction pathway acting in photoreceptor cells were pronounced (6 of 35 proteins described in KEGG pathway ola04744, false discovery rate 3.00*10^−1^, Fig3 G). For the upregulated ones, the eye samples showed the same trend as the whole organism, i.e. upregulation of the glycosylation machinery involving protein processing in the ER (5 of 155 proteins described in KEGG pathway ola04141, false discovery rate 2.46*10^−1^) as well as nucleotide sugar metabolism (Fig 3H).

In addition to the repeated upregulation of Gale, these were the Glutamine--fructose-6-phosphate transaminase 1 (Gfpt1) that catalyzes the formation of Glucosamine-6-phosphate which is a precursor for UDP-GlcNAc (Oki *et al*, 1999), the Dolichyl-phosphate beta-glucosyltransferase (Alg5) that provides the Dol-P-Glc and the Phosphomannomutase 2 (Pmm2, p = 0.087) that catalyzes the conversion of Man-6-P to Man-1-P as a precursor for Dol-P-Man. The ER-resident enzymes involved in core *N*-glycan processing to trim the glucoses in the α1,3-arm were upregulated as well (Zuber *et al*, 2000). The first Glucose removal is catalyzed by Mannosyl-Oligosaccharide Glucosidase (Mogs), followed by the Glucosidase II Alpha Subunit (Ganab) and Protein Kinase C Substrate 80K-H (Prkcsh) that trim the next two glucoses (Grinna & Robbins, 1979; Burns & Touster, 1982; Grinna & Robbins, 1980). Members of the subsequent protein folding control (Hspa5 and Hsp90b1) as well as the ER-to-Golgi transport system were upregulated in addition (Lman1, Sec23a, Sec22bb; (Khoriaty *et al*, 2012)).

Taken together, the proteomics analysis pointed on the one hand towards a compensatory upregulation of parts of the *N*-glycosylation pathway (Fig 3H). On the other hand, it revealed a massively affected phototransduction machinery (Fig 3G) with potential impact on the function of the retina that we have further addressed in detail.

### *Retinitis pigmentosa* caused by *alg*^*2hypo*^-mediated hypo-*N*-glycosylation

To investigate the state of neuronal structures like the brain and eyes, we performed Hematoxylin and eosin (H&E) staining on transverse sections of the head of stage 40 *alg2^hypo^* mutant- and wt-hatchlings. This revealed neuronal abnormalities reflected by reduced white matter in the mid- and hindbrain as well as in the optic tectum of *alg2^hypo^* hatchlings (Fig EV3), validating the phenotype described for ALG2 index patients in our fish model. Also the eyes of the mutants were slightly smaller in diameter compared to wt siblings (Fig 4A). Although the mutant retinae were apparently laminated, the inner plexiform layer (IPL) was not as pronounced as in wt siblings and the outer nuclear layer (ONL) was thinner by approximately one row of cells (Fig 4A’). In addition, the outer segments of the photoreceptors that usually extend into the retinal pigmented epithelium (RPE) were shortened. Since the ONL is a bilayer composed of rod and cone photoreceptors, we addressed its composition by immunofluorescence using the rod cell marker rhodopsin and the cone cell marker Zpr1 (Fig 4B). While the cone photoreceptors were not affected as highlighted by unaltered Zpr1 staining (Fig 4B upper panels), rod cells were absent from the *alg2^hypo^* mutant retinae as revealed by the rhodopsin antibody staining. The wt counterparts, in contrast, showed staining along the entire ONL. Since in *alg2^hypo^* mutant retinae only the rod photoreceptors were missing, we asked whether this was caused by a failure in rod photoreceptor differentiation or maintenance.

**Figure 4.**
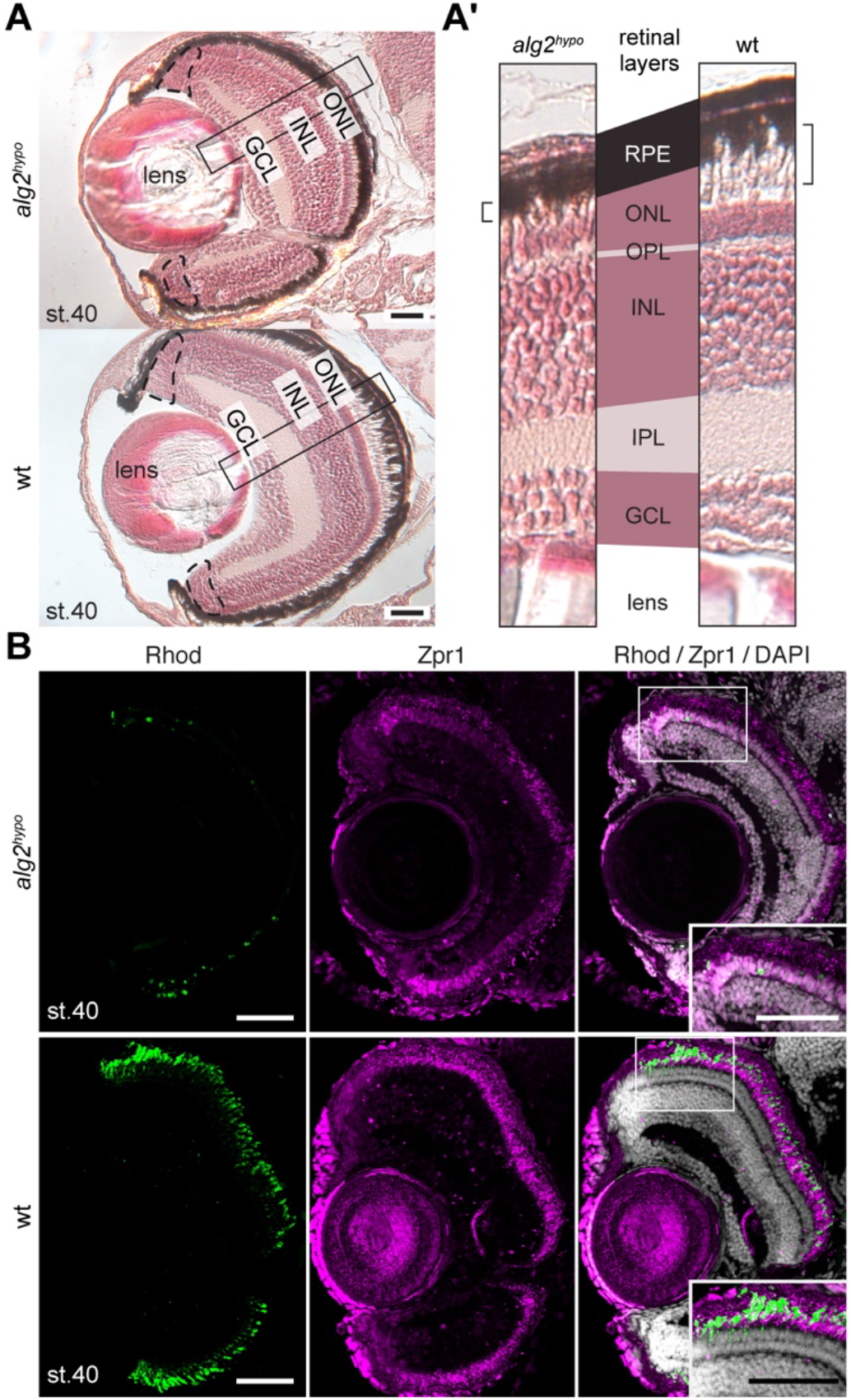
Rod photoreceptors are specifically absent from *alg2^hypo^* mutant retinae. Hematoxylin and eosin staining on transverse sections of *alg2^hypo^* homozygotes (upper panel) and wild type (wt) embryos (lower panel) at stage 40 (A, A’). A) Eyes of the *alg2^hypo^* hatchlings developed all retinal layers seen in wt. Peripheral stem cell niche of the eye indicated by dashed line. A’) Comparison of central retinal areas from sections in (A). Note: all retinal layers are comparable in size except for the outer nuclear layer (ONL) being reduced by the size of one row of cells and the inner plexiform layer (IPL) being slightly thinner. Note: photoreceptor outer segments (brackets) did not extend into the retinal pigmented epithelium (RPE) in *alg2^hypo^* mutants. B) Immunohistochemistry staining against photoreceptor specific markers rhodopsin (Rhod, rod photoreceptors, green) and Zpr1 (cone photoreceptors, magenta) and DAPI staining (grey) of stage 40 *alg2^hypo^* (upper panels) and wt (lower panels) transverse cryosections through the eye. Note: in contrast to wt, Rhod staining in mutants was restricted to peripheral areas of the retina (zoom in inset) whereas Zpr1 labels cone photoreceptors lining the entire ONL. GCL, ganglion cell layer; INL, inner nuclear layer; OPL, outer plexiform layer; scale bars = 50 μm

While stem and progenitor cells that reside in the ciliary marginal zone (CMZ) build to the growing retina (including all photoreceptors, (Centanin *et al*, 2014)), Müller Glial (MG) cells are triggered to divide and compensate for the acute loss of photoreceptors (Lust & Wittbrodt, 2018). We employed Rx2 as a bonafide marker for photoreceptors, stem and progenitor cells in the CMZ as well as MG cells (Reinhardt *et al*, 2015). Rx2 staining underpinned the absence of a layer of photoreceptor cells in the mutant compared to the photoreceptor bi-layer in wt hatchlings (Fig 5A, inset), consistent with histological analysis. The shape of the CMZ and number of MG cells, in contrast, were indistinguishable between *alg2^hypo^* and wt siblings, indicating proper proliferation and differentiation of the growing retina.

**Figure 5.**
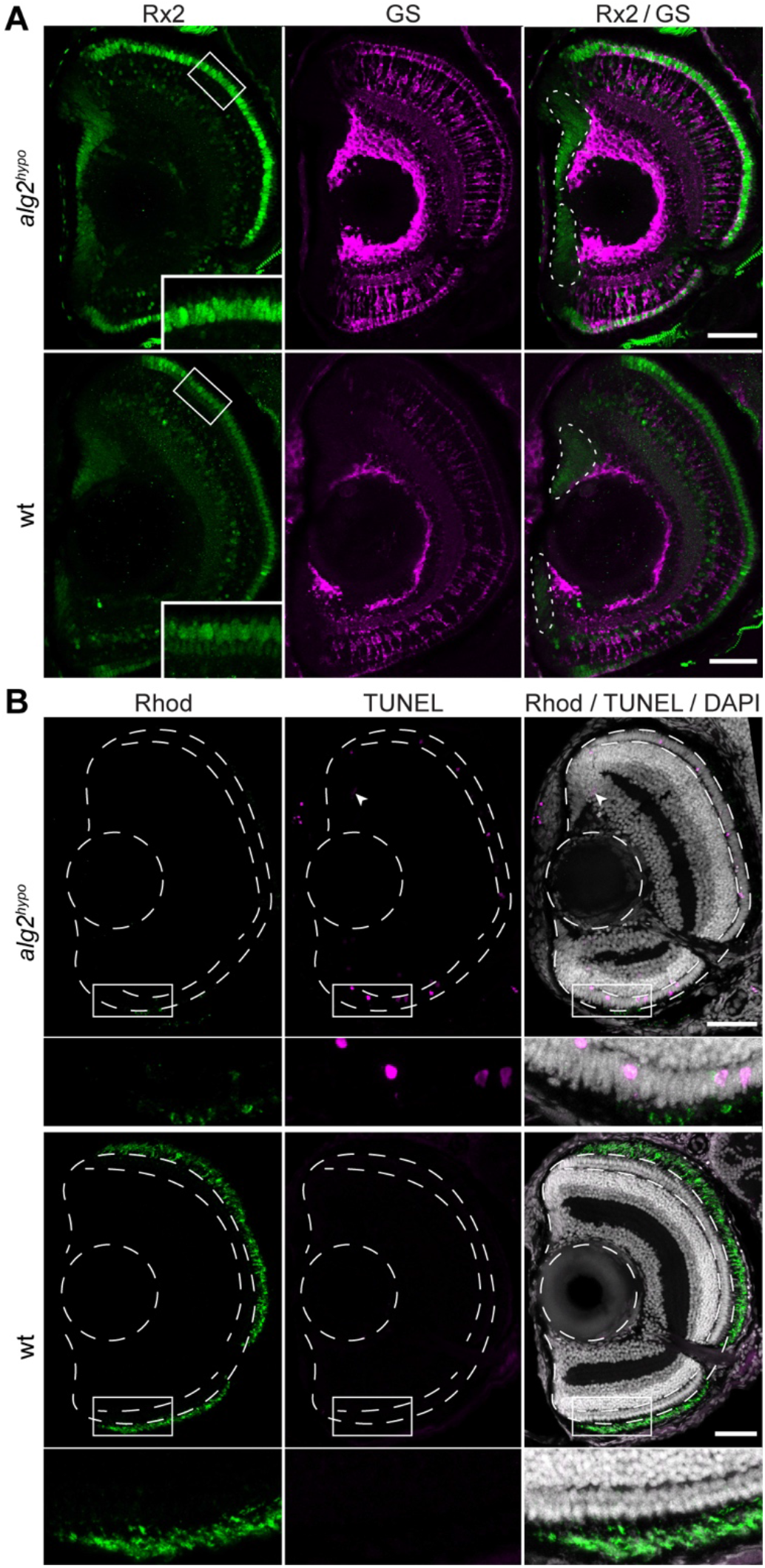
Rod photoreceptor specific apoptosis causes retinitis pigmentosa phenotype in *alg2^hypo^* mutants. Transverse cryosections through the eyes of stage 40 *alg2^hypo^* and wild type (wt) hatchlings. A) Immunohistochemistry against Rx2 (green) as a marker for retinal stem and progenitor cells, Müller-glia (MG) cells and photoreceptors and Glutamine synthetase (GS, magenta) highlighting MG cell bodies did not reveal alterations between *alg2^hypo^* (top panels) and wt controls (lower panels) in the CMZ or the central retina except for the outer nuclear layer. Here, Rx2 staining reconfirmed the monolayer of photoreceptors seen in *alg2^hypo^* hatchlings in contrast to the wt bi-layer (insets). B) TUNEL-positive cells are exclusively detected in the photoreceptor layer as well as in the progenitor zone of the stem cell niche (arrowheads) in *alg2^hypo^* homozygotes, but are not found in stage-matched wt samples (lower panels). Active loss of rod photoreceptors is reminiscent of a condition known as retinitis pigmentosa in patients. Scale bar = 50 μm

To address whether rod photoreceptor cells in *alg2^hypo^* mutants were actively eliminated, (programmed) cell death was visualized by TUNEL staining. Apoptotic cells were indeed especially enriched in the photoreceptor layer as well as in a subset of progenitor cells in the central part of the CMZ (Fig 5B, upper panels). Stage-matched wt siblings did not show apoptotic cells in the retina (Fig 5B, lower panels). This finding indicates that the rod photoreceptors were initially formed during development and post-embryonic growth, but were not maintained and were rather eliminated by apoptosis. This loss of rod photoreceptors is described as retinitis pigmentosa or night blindness.

### *Alg*^*2hypo*^ mutant can be rescued by exogenous supply of full length *alg2* mRNAs

To address the rescue potential of *Alg2* mRNA we microinjected full-length human or medaka *alg2* mRNA (100 −200 ng/μl) at the one cell stage. Injection of either mRNA efficiently rescued the gross morphology phenotype and expanded the life-span by at least 16 days. Gross morphology was fully restored in injected *alg2^hypo^* mutant embryos (medaka *alg2* n = 11, human *Alg2* n=2; genotypically confirmed homozygotes) while control injection with *GFP* mRNA (100 −200 ng/μl) had no rescue effect (n = 4) (Fig 6A). The exogenous supply of medaka *alg2* mRNA did neither lead to apparent phenotypes in heterozygous *alg2^hypo^* (n = 22) nor wt siblings (n = 15). Alcian blue staining of injected embryos (Fig 6B) showed a full reversion of the head cartilage phenotype in all rescued *alg2^hypo^* mutant embryos (p = 0.007, p = 0.02, p = 0.11 for Meckel’s cartilage, ceratohyal, and palatoquadrate cartilages, respectively; two-tailed nonparametric Student’s T-test; Fig 6C). Cartilage lengths were indistinguishable between the rescue injections and medaka *alg2* overexpression in wt embryos, respectively (p = 0.48, p = 0.97, p = 0.61 for Meckel’s cartilage, ceratohyal, and palatoquadrate cartilages, respectively; two-tailed nonparametric Student’s T-test; Fig 6C). Injection of medaka *alg2* mRNA not only rescued gross morphology, but also restored viability (at least) until the fish were sacrificed for downstream analysis at 18 dph, while *GFP* control injected or uninjected *alg2^hypo^* hatchlings died at 2-3 dph. The apparent problem of *alg2^hypo^* hatchlings to maintain rod photoreceptors detailed above was efficiently overcome upon the mRNA mediated rescue. Analysis of rod- and cone-photoreceptors by immunohistochemistry on retinal cross-sections demonstrated the full restoration of the bilayered structure of the ONL in the entire lining of the retina of rescued *alg2^hypo^* hatchlings, which was indistinguishable from wt controls (Fig 4B) or wt siblings overexpressing medaka *alg2* mRNA (Fig 6D).

**Figure 6.**
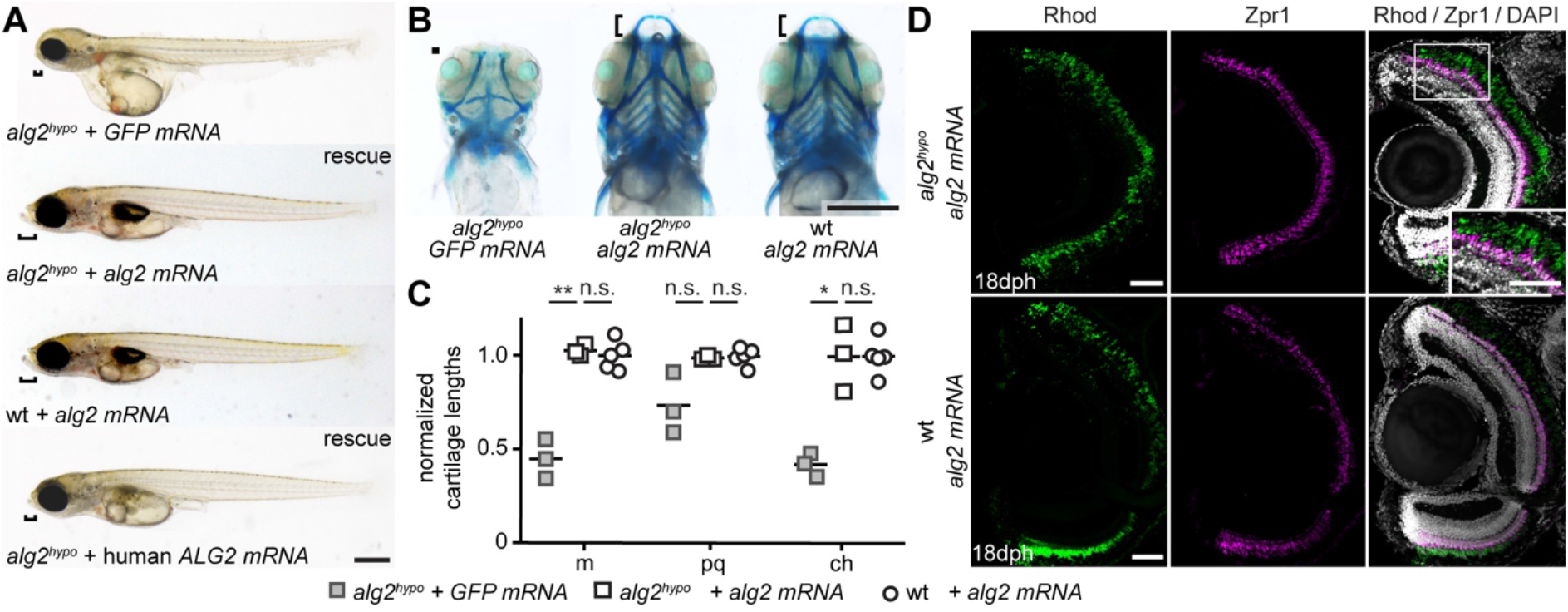
Human and medaka *alg2* mRNA rescue multisystemic phenotype in *alg2^hypo^*. A) The multisystemic phenotype seen in *alg2^hypo^* hatchlings can be fully rescued with full length medaka *alg2* mRNA but not *GFP* mRNA control injections at the one cell stage. Exogenous supply of *alg2* mRNA in wildtype (wt) siblings did not provoke any phenotype. Also full length human *ALG2* mRNA rescued the gross morphology phenotype, yet to a lesser extent evident in partially rescued snout size (brackets). B) Alcian blue staining (ventral view, stage 40) of *GFP* mRNA control (left) and full length medaka *alg2* mRNA rescue (central) injected at the one cell stage into *alg2^hypo^* zygotes as well as full length medaka *alg2* mRNA injected wt (right) specimens. Cartilage anatomy in rescued and exogenous supplied *alg2* in wt was indistinguishable. C) Quantitative analysis of cartilage lengths, normalized to standard length and mean of wt (cf. Fig 1F). Full length *alg2* mRNA completely rescues craniofacial cartilage structures whereas control *GFP* mRNA injections did not. Exogenous *alg2* mRNA does not alter cartilage structures. m, Meckel’s cartilage; pq, palatoquadrate; ch, ceratohyal. Scale bars = 0.5 mm.

Taken together, the medaka *alg2^hypo^* line presented in this study is a fully rescuable organismal model for hypo-*N*-glycosylation, recapitulating several clinical hallmarks of CDG-I deficiencies. The discovery of misregulated retina specific proteins and the apparent loss of rod photoreceptors in the model hint at retinitis pigmentosa as an additional, so far unnoticed human phenotype for ALG2-CDG.

## Discussion

Studying the mechanism and consequences of protein glycosylation had been very challenging due to the crucial importance of the pathway for protein function and consequently cellular and organismal survival. A failure to properly glycosylate in human patients results in CDGs, rare genetic diseases (Chang *et al*, 2018). The unique genetic variants in key glycosylation factors carried by the patients provide essential information on the structure-function relationship of the proteins involved. To address the function and regulatory interplay of the complex glycosylation network, we have established an Alg2 animal model and investigated the molecular, cellular and morphological consequences of a particular mutation modeled in analogy to the allele of an ALG2 index patient. We have uncovered a striking compensation on the level of the glycosylation network revealing a complex regulatory interplay.

For both the medaka mutant hatchlings and the Alg2 index patient fibroblasts, we detected a distinct hypo-*N*-glycosylation that was less pronounced in the case of the patient fibroblasts. To study the molecular and cellular consequences of lowering the abundance of N-glycan structures to a critical minimum, we further employed our *alg2^hypo^* animal model. It represents a sensitized situation that facilitated the identification of proteins and pathways that crucially depend on proper glycosylation. Comparing wt and mutant animals and organs respectively, we observed two prominent consequences: on the one hand the compensatory upregulation of the glycosylation machinery and on the other hand the specific downregulation of proteins involved in phototransduction resulting in the loss of a single cell type in the retina.

On the side of compensatory effects, our proteomics results pointed at two routes by which the affected organism counteracted the reduced activity of the *alg2^hypo^* variant. We detected an upregulation of enzymes involved in the nucleotide sugar metabolism, resulting in higher levels of the enzymes that deliver the glycan building blocks (Gale, Gfpt1, Alg5, Pmm2). On the other hand we detected an upregulation of the machinery that processes the core *N*-glycan inside the ER-lumen, acting downstream of Alg2 (Fig 7).

**Figure 7.**
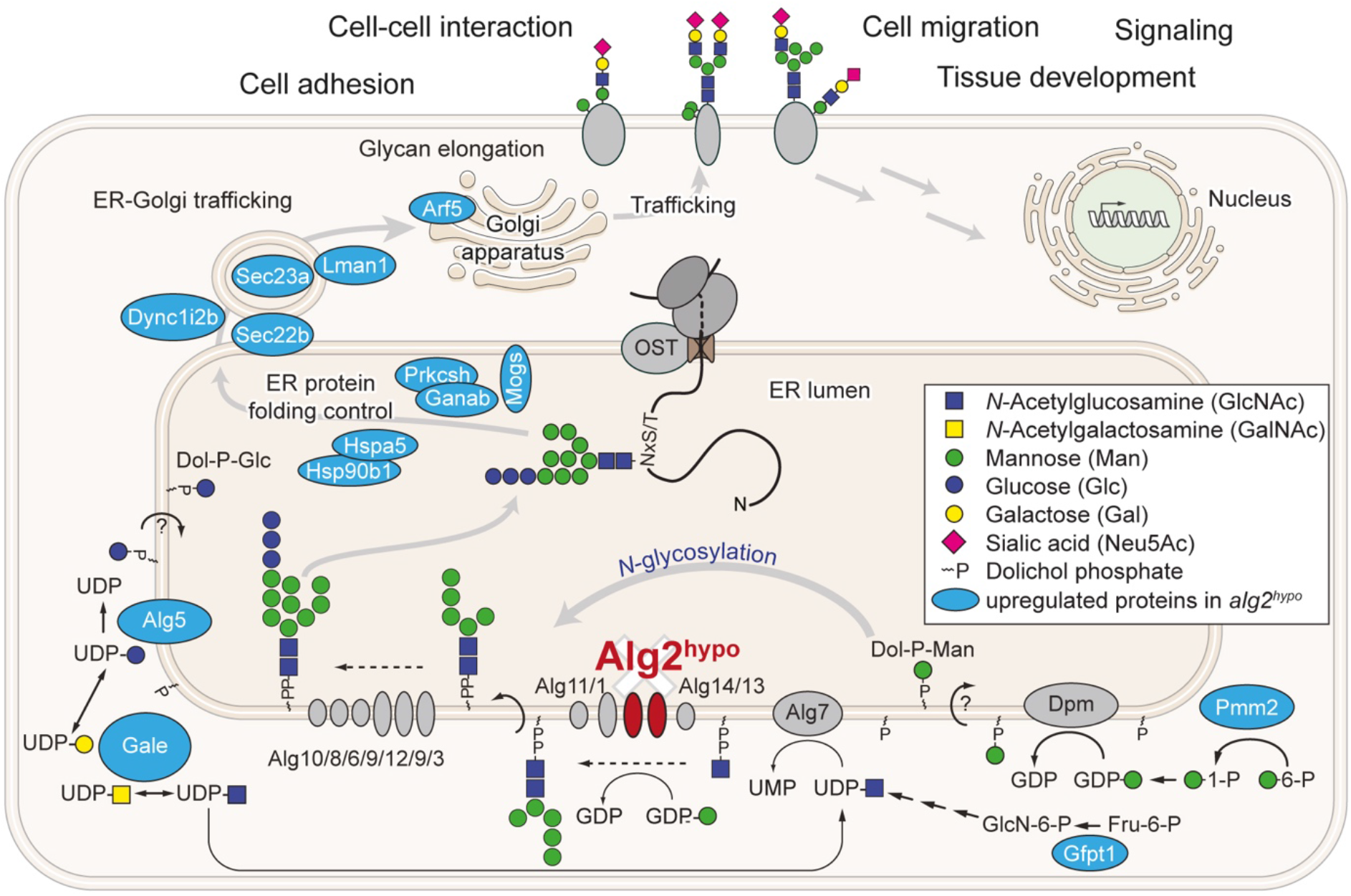
The upregulation of the glycosylation machinery as compensatory effort in *alg2^hypo^* mutants. Schematic overview of altered *N*-glycosylation machinery detected in *alg2^hypo^* mutants with the upregulated proteins found in the proteomics analysis (light blue) put into context. Two main routes affecting glycosylation can be seen: upregulation of basal enzymes allocating nucleotide sugars (Gale, Gfpt1, Alg5 and Pmm2) and following the translocation of the oligosaccharide from the lipid-linked donor to the Asn of a nascent protein, factors relevant for core *N*-glycan processing (Mogs, Ganab, Prkcsh), ER resident protein folding control (Hspa5, Hsp90b1) and ER-to-Golgi trafficking (Sec22b, Dyn1l2, Sec23a, Lman1, Arf5) were highly abundant. OST, Oligosaccharyltransferase

Providing more nucleotide sugars for the glycan synthesis can be regarded as compensatory action to facilitate the synthesis of sufficient levels of the core *N*-glycan under conditions of reduced Alg2 activity. The xCGE-LIF analysis showed reduced levels of known *N*-glycans, while the structure of the *N*-glycans transferred onto proteins was unaffected. It can thus be concluded that only properly generated, yet rare Glc3Man9GlcNAc2 was translocated from the dolichol donor-LLO to an Asn at the glycosylation site of nascent proteins. To prime the cell for the subsequent *N*-glycan processing, enrichment of the downstream machinery was evident: the three Glucoses in the alpha-1,3-arm of the core *N*-glycan have to be trimmed not only as checkpoint control for proper protein folding, but also to ensure that processing to mature oligosaccharides can continue (Zuber *et al*, 2000). All three enzymes (Mogs, Ganab, Prkcsh) involved in this process were highly upregulated in the eye samples of the *alg2^hypo^* variant. On the route of the *N*-glycosylated proteins towards full oligosaccharide processing, the newly generated glycoprotein is shuttled via vesicle transport from the ER to the Golgi, involving the upregulated Sec22b, Sec23a and Lman1 as well as Arf5 and Dync1l2. Although upregulated, the compensatory effort did not suffice for the full maintenance of all cells and tissues in the *alg2^hypo^* mutants. Our analysis revealed the framework of adjustments an organism compensating hypo-*N*-glycosylation employs, providing a glimpse on the closely intertwined regulatory topology of the glycosylation network.

Besides the upregulation of the glycosylation machinery, we detected an enrichment of the master regulator for ER stress, Hspa5 (Lee, 2005) and its interactor Hsp90b1 both part of a large ER-localized multiprotein complex for protein folding control (Meunier *et al*, 2002). We interpret this as a response to the higher abundance of proteins failing to pass the ER-quality checkpoint due to improper folding in the absence of *N*-glycosylation. Eventually, prolonged ER stress will induce apoptosis (Szegezdi *et al*, 2006).

The second prominent group of ALG2 dependent proteins revealed by our proteomics analysis are photoreceptor specific and involved in phototransduction as confirmed via immunofluorescence. Since retinal phenotypes have been reported for several CDGs, we have analyzed retinal histology and cell type composition in detail and have uncovered the progressive, specific loss of rod photoreceptors, a hallmark of retinitis pigmentosa (RP). RP can be caused by several different inherited mutations all resulting in a progressive loss of vision as a result of abnormalities in rod and cone photoreceptors or the RPE (Ferrari *et al*, 2011). We have shown that in *alg2^hypo^* mutants rod photoreceptors are initially born, but cannot be maintained. Consequently, the underrepresentation of retinal proteins in the mutant eyes can be interpreted as a direct consequence of the loss of cells. Alternatively, downregulated proteins could represent direct Alg2 targets that require glycosylation for rod photoreceptor survival. Strikingly, three of the prominently underrepresented retinal proteins Rhodopsin, Arrestin and a violet sensitive opsin (Rho, Kfharr2 and Opn1sw) contain glycosylation sites, even though not a signal peptide by prediction (NetNGlyc 1.0, (Gupta *et al*)). For Rhodopsin however, it has been demonstrated that its function and the survival of photoreceptors crucially depend on N-glycosylation of conserved asparagines at the N-terminus (Asn-2 and Asn-15) (Murray *et al*, 2009). Targeted mutation of Asn-15 leads to defects in Rhodopsin folding and its transport to the cell surface, eventually resulting in prolonged ER stress and rod cell specific death (Kaushal *et al*, 1994; Tam & Moritz, 2009).

It is intriguing that the *alg2^hypo^* variant specifically triggers apoptosis of the rod cells, while none of the other neuronal cell types in the eye were apoptotic. This highlights that some cell types crucially depend on *N*-glycosylation of specific target proteins and consequently, their maintenance critically relies on the continuous supply of sufficient levels of *N*-glycans. This was only detectable in the organismal context and underpins the high relevance of model systems for providing the full perspective when studying the multisystemic, cell-type specific impact of glycosylation and its absence, respectively. Even though Alg2 is expressed ubiquitously, crucial targets are cell-type specific and might not be expressed at all if tissue culture models are employed.

The complex, multisystemic phenotype of the *alg2^hypo^* mutant that becomes apparent only from stage 39 onwards was efficiently rescued by injection of *alg2* mRNA at the one-cell stage. This surprising finding indicates that the early production of wt Alg2 protein within the first 24 hours sufficed to rescue the multisystemic phenotypes. Rescued *alg2^hypo^* mutants were fully viable until they were sacrificed for histological analysis at juvenile stages and were indistinguishable from their heterozygotic and wt siblings. Given the limited stability of injected mRNA, the rescue of embryonic phenotypes and survival up to experimental termination at juvenile stages is consistent with a transient requirement of Alg2 activity during early development or a slow turnover rate of the Alg2 protein or a combination of both.

The high and long lasting rescue potential of mRNA injected into *alg2^hypo^* mutants could be indicative of a transient requirement of ALG2 activity during embryonic development. Once hypo-*N*-Glycosylation is overcome by the rescuing mRNA, the resulting glycosylation levels would suffice for tissue maintenance. Interestingly, ALG enzymes, and in particular ALG2, are very low abundance proteins in resting cells with a half-life of 240 days in HeLa (Zecha *et al*, 2018). Its stability may even be increased in complexes with ALG1 and ALG11 (Gao *et al*, 2004). Consequently a transient pulse of exogenous wt *alg2* mRNA establishes a uniformly distributed, stabilized, long lasting pool of rescuing Alg2 protein. The apparent transient requirement may thus be best explained by a long lasting effect of protein stability. The *alg2^hypo^* mutant model described here offers the unique opportunity to tackle linear structure-function relationships by rescue experiments with *in vitro* modified mRNA variants. Future analyses combining mRNA based rescue strategies with acute protein depletion will pave the way for integrating the structure-function relationships with the delineation of cell type specific requirements of Alg2.

## Materials and Methods

### Animal husbandry and ethics statement

Medaka (*Oryzias latipes*) Cab strain used in this study were kept as closed stocks in accordance to Tierschutzgesetz §11, Abs. 1, Nr. 1 and with European Union animal welfare guidelines. Fish were maintained in a constant recirculating system at 28°C on a 14 h light/10 h dark cycle (Zucht-und Haltungserlaubnis AZ35-9185.64/BH).

Written informed consent was obtained for analysis of patient-derived material. This study was approved by the Ethics Committee of the Medical Faculty Heidelberg.

### designing and cloning of sgRNAs

sgRNAs were designed with CCTop and standard conditions (Stemmer *et al*, 2015). The following target sites were used (PAM in brackets): *alg2 T1* (5-CCCGTTATTGCCGTCAACTC[TGG]), *alg2 T2* (5-CCGTTATTGCCGTCAACTCT[GGG]). Cloning of sgRNA templates and *in vitro* transcription was performed as detailed in (Stemmer *et al*, 2015).

### Generation of animal lines and genotyping

One-cell stage wt medaka (*Oryzias latipes*) Cab strain zygotes were microinjected into the cytoplasm with 150 ng/μl *Cas9* mRNA, 15 ng/μl per sgRNA, 10 ng/μl single-stranded oligodeoxynucleotide (ssODN) that introduces a premature STOP codon (underlined) upon proper integration 5-GGTCTCTGATGAGCCTCTCCATGGCCTGGGAGAACGCCTCAGCCGTAGGCTCGCACA GGAAGCCCGTCTCCCCGTGTGCTACGCTCTCCAGAGGGCC**<u>TTA</u>**AGAGTTGACGGCAAT AACGGG-3 and 10 ng/μl *GFP* mRNA as injection tracer. GFP positive crispants were raised, outcrossed to wt and genotyped from extracted DNA of fin clip biopsies in extraction buffer (0.4 M Tris-HCl pH 8.0, 5 mM EDTA pH 8.0, 0.15 M NaCl, 0.1% SDS in distilled water with 1mg/ml Proteinase K (Roche, 20 mg/ml)). Genotyping-PCR was performed with Q5^®^ High-Fidelity DNA Polymerase (New England Biolabs) with 98 °C initial denaturation for 2 min, followed by 30 cycles of: 98 °C denaturation for 20 sec, 67 °C annealing for 30 sec, 72 °C extension for 25 sec. Primers used: forward 5-TCCACTTGGAGGATTGCGTC, reverse 5-CATTTAGCTGGGGATTGGTACAC. Heterozygosity was assessed with T7 endonuclease I cleavage (New England Biolabs, #M0302S) upon direct incubation of 10 μl amplicon, 2 μl 10x NEB buffer 2, 7.5 μl ddH_2_O and 0.5 μl T7E1 (NEB) for 30 min at 37°C. Fragments were size separated and analyzed via gel electrophoresis. Knock in of the ssODN donor was assessed with StuI test digest of the genotyping amplicon using primers described above, at 37 °C for 2 hours following agarose gel electrophoresis and verified by sequencing (MWG eurofins).

Embryos were maintained in ERM (170 mM NaCl, 4 mM KCl, 2.7 mM CaCl_2_.2H_2_O, 6.6 mM MgSO_4_.7H_2_O, 170 mM HEPES) with 0.00002% (w/v) methylene blue at 26-28 °C upon collection from female adults.

### Fibroblast culture maintenance

Patient and control fibroblasts were cultured in Dulbecco’s modified Eagle’s medium (high glucose; Life Technologies) supplemented with 1 % FCS (PAN Biotech) and 1 % Pen/Strep under 5 % CO_2_ at 37 °C. Medium was replaced every 72 h.

### -Alcian blue staining

Stage 40 (9 dpf at 26 °C) medaka hatchlings were anaesthetized with 1X Tricane, tail was clipped and processed for PCR-genotyping. Specimens were fixed in 4% PFA in PBS overnight at 4 °C. Samples were dehydrated in 50% and 70% EtOH for 15 min at RT. Cartilages were stained in 0.02 % Alcian Blue 8GX (Sigma Aldrich) in 70 % EtOH and 60 mM MgCl_2_ overnight at RT. Hatchlings were washed once in 70% and 50% EtOH and eye pigments were bleached in 1 % KOH, 3 % H_2_O_2_ in PBS for 30 min at RT. Hatchlings were washed once with 50% EtOH and imaged under Nikon SMZ18 in 3 % methylcellulose. For long term storage, samples were dehydrated in 100 % EtOH and kept at −20°C. Using Fiji (Schindelin *et al*, 2012), analysis of cartilage lengths (euclidean distance) was performed according to (Cline *et al*, 2012) and two-tailed nonparametric Student’s T-test was applied for statistics.

### Hematoxylin and eosin staining

Stage 40 (9 dpf at 26°C) medaka hatchlings were fixed in Davidson’s fixative (3 parts tap water, 3 parts EtOH, 2 parts formalin, 1 part 98 % acetic acid) overnight at 4 °C. Samples were dehydrated in 70 % −90 % −100 % EtOH and xylene series. Dehydrated hatchlings were incubated in paraffin for 1 h at 60 °C. All samples were embedded in paraffin blocks at RT and sectioned on microtome with 7 μm thickness and mounted on glass slides. Following overnight incubation at 42 °C, samples were stepwise rehydrated in xylene – 100 % −90 % −70 % −50 % EtOH series. Slides were stained in hematoxylin for 10 min, washed with tap water for 40 min and stained with eosin for 2 min at RT. Samples were dehydrated with 70 % −90 % −100 % EtOH and xylene series. Samples were preserved with Eukitt® Quick-hardening mounting medium (Sigma-Aldrich). Images were taken under DIC microscope (Leica DB5000).

### Antibody staining on cryosections

Samples were fixed in 4 % PFA in PTW (PBS-, 0.4 % Tween) overnight at 4°C. Samples were washed in PTW and incubated once overnight in 30 % sucrose in PTW and once in Tissue Freezing Medium (Leica, #14020108926): 30 % Sucrose in PTW (1:1, *w*/*w*). Samples were embedded in the same Tissue Freezing Medium and sectioned with 16 μm thickness on a cryostat (Leica CM 3050S). Slides were incubated overnight at 4 °C and washed with PTW for rehydration. Slides were blocked with 10 % normal goat serum (NGS) in PTW for 2 h at RT. Samples were stained with anti-Rhodopsin rabbit and mouse antibodies (1:200, home-made and 1:200, Millipore, #MABN15), anti-Zpr1 mouse antibody (1:200, ZIRC, #AB_10013803), anti-Rx2 rabbit antibody (1:200, (Reinhardt *et al*, 2015)), and anti-GS mouse antibody (1:500, Merck #MAB302) in 1 % NGS overnight at 4 °C. Goat anti-mouse IgG (H+L) Alexa Fluor 546 (Life Technologies, #A11030) and goat anti-rabbit IgG Alexa Fluor 488 (Life Technologies, #A11034), and donkey anti-mouse 647 (Invitrogen, #A32787) were used as secondary antibodies together with DAPI (10 μg/ml) in 1 % NGS for 2 h at 37 °C. When combined with TUNEL staining, *In Situ* Cell Death Detection Kit, TMR red (Roche, #12156792910) was used according to the manufacturer’s instruction. Samples were mounted in 60 % glycerol in PTW and were imaged on a Leica SP8 confocal microscope. Image analysis was performed with Fiji (Schindelin *et al*, 2012).

### Lectin blots

#### Medaka

Total proteins were isolated from pool of 10-20 hatchlings (stage 40, anaesthetized) with RIPA Lysis and Extraction Buffer (Thermo Scientific, #89900) including cOmplete™ EDTA-free Protease Inhibitor Cocktail (Roche, #1183617001). Quantification performed with Pierce™ BCA Protein Assay Kit (#23225). 10 μg of protein was loaded into each lane of 10 % SDS-PAGE. Proteins were blotted on PVDF membrane (Millipore Immobilon-P, #IPVH00010) at 100 V for 1 h at 4 °C. Membrane was blocked with 5 % BSA in 1X TBST (TBS with 0.1 % Tween) for 1 h at RT. As an internal control, anti-Gapdh rabbit monoclonal antibody (Cell Signaling(14C10), #2118) was used in 1:1000 dilution in blocking buffer (5 % BSA in 1X TBST) for 1 h at RT. Goat anti-rabbit HRP (Agrisera, #AS09602) was used in 1:5000 concentration as a secondary antibody. After developing signal with Pierce™ ECL Western Blotting Substrate (Thermo Scientific, #32109), the blot was stripped in mild stripping buffer (1.5 % glycine (*w*/*v*), 0.1 % SDS (*w*/*v*), 1 % Tween20 (*w*/*w*), pH 2.2). Blots were incubated in streptavidin solution (1 drop in 10 ml TBST, Vector laboratories #SP-2002) 15 min at RT to block internal biotin signal of the fish. Blots were incubated with either biotinylated Concanavalin A (Con A, Vector Laboratories, #B-1005) or biotinylated Wheat Germ Agglutinin (WGA, Vector Laboratories, #B-1025) at 1:1000 dilution in TBST 2.5 hr at RT. Streptavidin-Horseradish Peroxidase (Vector Laboratories, #SA5004) was used at 1:10000 dilution for 30 min at RT. Signal was developed with Pierce™ ECL Western Blotting Substrate (Thermo Scientific, #32109).

#### Fibroblast

Cell monolayers were washed with ice-cold PBS and harvested by cell scraper. Cells were lysed 30 min in RIPA buffer (ThermoFisher Scientific) on ice and by passing the sample 20 times through a 20G needle. Samples were centrifuged for 30 min at 13,000 rpm at 4 °C. 10 μg of total protein derived from patient and control fibroblasts were used for loading. Samples were mixed with 6x Laemmli buffer (375 mM Tris–HCl, pH 6.8, 6 % SDS, 48 % glycerol, 9 % 2-mercaptoethanol, 0.03 % bromophenol blue) and denatured at 95 °C for 5 min. Extracts were analyzed on a 12.5 % SDS-PAGE and blotted onto a nitrocellulose membrane (GE Healthcare) by semi-dry electrophoretic transfer. The membrane was blocked for 1 h at room temperature with 5 % milk powder in PBST (0.1 % Tween20 in PBS). After blocking, the membrane was washed and incubated with the primary antibodies against ß-actin antibody (1:10,000, Sigma #A5441) overnight at 4 °C. After washing, the membrane was incubated with secondary antibody anti-mouse IgG-conjugated with horseradish peroxidase (Santa Cruz, Dallas, USA; 1:10,000) for 45 min at RT. Protein signals were detected by light emission with Pierce™ enhanced chemiluminescence reagent (ECL) plus western blot analysis substrate (ThermoFisher Scientific). After stripping the blots with 10 % acetic acid for 12 min the membrane was blocked with 5 % BSA in TBST (0.5 % Tween20 in TBS) for 1 h. Membranes were incubated with the biotinylated lectins ConA and WGA (Vector Laboratories) at 1:1000 dilution in TBST for 2 h at RT. The membranes were subsequently incubated with horseradish peroxidase streptavidin (Vector Laboratories) for 30 min and detected with PierceECL plus assay kit (ThermoFisher Scientific).

### Multiplexed capillary gel electrophoresis with laser induced fluorescence (xCGE-LIF)

Sample preparation and quantitative analysis by xCGE-LIF was performed according to a modified version of previously described protocols (Thiesler *et al*, 2016; Hennig *et al*, 2016). Briefly, samples from pools of 20 medaka hatchlings (stage 40) and human fibroblasts (7×10^5^ cells) were lysed with RIPA Lysis and Extraction Buffer (Thermo Scientific, #89900). Samples were purified with methanol/chloroform protein extraction protocol according to (Wessel & Flügge, 1984). The following sample preparation was carried out with the glyXprep 16 kit (glyXera, Magdeburg, Germany). *N*-glycans were released from solubilized proteins using peptide-*N*-glycosidase F and fluorescently labeled with 8-aminopyrene-1,3,6-trisulfonic acid (APTS). Excessive fluorescent dye was removed by hydrophilic interaction liquid chromatography-solid phase extraction (HILIC-SPE). The purified APTS-labeled *N*-glycans were analyzed by xCGE-LIF. Data processing and normalization of migration times to an internal standard were performed with glyXtool™ software (glyXera, Magdeburg, Germany). *N*-glycan fingerprints (normalized electropherograms) were annotated based on migration time matching with an in-house *N*-glycan database (glyXbase™) and exoglycosidase sequencing. The symbolic representations were drawn with GlycoWorkbench (Ceroni *et al*, 2008) according to the guideline of the Consortium for Functional Glycomics (Varki *et al*, 2009). To facilitate the quantitative inter-sample comparison, an aliquot of each sample was spiked in with 1 μg of a bovine asialofetuin as an internal standard prior to methanol/chloroform protein extraction. A unique and asialofetuin-derived *N*-glycan peak was used to quantitatively normalize peak intensities. The quantitative normalization was then applied to the standard *N*-glycan fingerprints using mannose-6 as a transfer peak as explained in Fig EV2.

### Full-length mRNA rescue

#### Cloning of Alg2 cDNA and mRNA synthesis

*Alg2* cDNA was amplified with RT-PCR from the cDNA of wt *Oryzias latipes* Cab strain stage 18 embryos with Q5® High-Fidelity DNA Polymerase (New England Biolabs) by using primers with BamHI-HF (New England Biolabs, 20U/ml, #R3136S) and XbaI (New England Biolabs, 20U/ml, #R0145S) recognition sequence extensions, 5-GCCGGATCCATGGCGCGGGTGGTGTTT-3 and 5-GCCTCTAGATTACTGGCTGAGCATAACTACGT-3, respectively (58°C annealing for 30 sec, 70 °C extension for 40 sec, 35 cycles). PCR product and pCS2+ vector (Rupp *et al*, 1994) were digested with BamHI-HF and XbaI restriction enzymes and cleaned up from agarose gel with innuPREP Gel Extraction Kit (AnalytikJena). Digested PCR product and backbone were ligated with 0.5 μl T4 DNA ligase (Thermo Scientific, 5U/ml) in 1x ligase buffer, 10 μl end volume for 15 min at RT. Cloned vector was transformed into Mach1-T1 cells (ThermoFisher Scientific) via with heat shock induction at 42 °C for 45 sec and snap cooling on ice, 300 μl TB added and incubated for 45 min at 37 °C. 100 μl were plated on LB plates with Ampicillin resistance for overnight incubation at 37 °C. Individual clones were inoculated into LB medium containing Ampicillin (100 μg/ml). Plasmids were extracted from bacteria culture with QIAprep Spin Miniprep Kit (Qiagen, #27104). One clone with a successful integration was used to perform *in vitro* mRNA synthesis with mMESSAGE mMACHINE™ SP6 Transcription Kit (ThermoFischer Scientific, #AM1340) upon NotI-HF (New England Biosciences, 20 U/ml, #R3189S) linearization and gel purification with innuPREP Gel Extraction Kit (Analytik Jena AG). Remaining of the plasmid was digested with 1 μl TURBO DNase (2 U/μl, ThermoFisher Scientific) for 15 min at 37 °C. RNA was cleaned up with RNeasy Mini Kit (Qiagen, 74104) and quality of RNA was confirmed with agarose gel electrophoresis and spectrophotometry (Nanodrop).

The plasmid containing the whole cDNA of healthy human was kindly provided by Christian Thiel (University Clinics, Heidelberg University, Germany). The RNA was synthesized *in vitro* with mMESSAGE mMACHINE™ T7 Kit (Ambion, #AM1344) on HpaI linearized and gel purified (InnuPrep, AnalytikJena) DNA as described by the manufacturers.

#### Injections into medaka

Adults heterozygous *Alg2^hypo^* medaka were crossed and offspring was injected at the one-cell stage with 100 −200 ng/μl medaka *alg2* or 100 ng/μl of human *Alg2* mRNA both together with *GFP* mRNA (10 ng/μl) as injection tracer. As injection control, offspring of the same crossing scheme was used for *GFP* mRNA injection only. Embryos were kept at 28 °C. GFP negative embryos were discarded. Genotyping of the embryos was performed from the fin clip biopsies with Q5® High-Fidelity DNA Polymerase (New England Biolabs) as stated above. The remainder sample was forwarded to the particular analytic procedures detailed above.

### Mass Spectrometry

#### Sample preparation and protein precipitation

Mass spectrometry (MS) was performed on whole deyolked stage 40 hatchlings as well as stage 40 eyes. For the whole organism, stage 40 hatchlings were euthanized with tricane, deyolked, pooled (n=3 biological replicates with n=6 each), snap frozen in liquid nitrogen and kept at −80 °C until lysis. For the measurement of eyes, both left and right eyes were dissected from 15 fish per biological replicate (n=4 biological replicates, 30 eyes each) and snap frozen in liquid nitrogen. Samples were lysed with 100-150 μl of RIPA Lysis and Extraction Buffer (Thermo Scientific, #89900) including cOmplete™ EDTA-free Protease Inhibitor Cocktail (Roche, #1183617001) with the help of Qiagen TissueRaptor II. Protein lysis was incubated on ice with 50 U of Benzonase Nuclease (Millipore, #E1014-5KU) for 20 min and at 37 °C for 5 min. Samples were then centrifuged at 12000 g for 10 min at 4 °C. Supernatant was taken into fresh tubes and protein concentration was assessed via Pierce™ BCA Protein Assay Kit (#23225). 100 ug and 30 ug of total protein for whole hatchling and eyes, respectively, were used for protein precipitation. Samples were precipitated with methanol / chloroform protocol according to (Wessel & Flügge, 1984).

#### In-solution digestion

For in-solution digestion, pellets of precipitated proteins were resuspended in 20 μL of urea buffer (8 M urea, ≥ 99.5 %, p.a., Carl Roth, 100 mM NaCl, ≥ 99.5 %, p.a., Applichem, in 50 mM triethylammonium bicarbonate (TEAB), pH 8.5, Sigma-Aldrich). Cysteine thiols were subsequently reduced and alkylated by adding Tris(2-carboxyethyl)phospin (Carl Roth) to a final concentration of 10 mM and 2-Chloroacetamide (≥ 98.0 %, Sigma-Aldrich) to a final concentration of 40 mM. The solution was incubated for 30 min at RT. Sample pre-digestion was performed with Lysyl Endopeptidase® (MS grade, Wako Chemicals), which was added in an enzyme:protein ratio of 1:40 (*w*/*w*) before the sample was incubated for 4 hours at 37 °C. After diluting the urea concentration to 2 M by adding 50 mM TEAB buffer, trypsin (MS grade, Thermo) was added in an enzyme:protein ratio of 1:100 (*w*/*w*) and incubated for 16 hours at 37 °C.

#### Dimethyl Labeling

Dimethyl duplex labeling was performed bound to C18 material according to standard protocol (Boersema *et al*, 2009). Briefly, digestion reaction was stopped by reducing the pH < 2 through the addition of trifluoroacetic acid (TFA, ≥ 99.0 %, Sigma-Aldrich) to a final concentration of 0.4 % (v/v) and centrifuged for 10 min at 2500 g at RT. Supernatants volume corresponding to 20 μg of total tryptic peptides per sample were loaded onto C18 StageTips (Rappsilber & Mann, 2007) containing 3 disks of Empore C18 material (3M). Prior to sample loading, StageTip material was successively equilibrated with 20 μL of methanol (MS grade, Carl Roth), followed by 20 μL of 50 % (*v*/*v*) acetonitrile (ACN, UPLC grade, Biosolve) in 0.1 % (*v*/*v*) TFA and by 20 μL of 0.1 % (*v*/*v*) TFA with centrifuging for 1 min at 1500 g after each equilibration step. Loaded peptide samples were washed with 20 μL of 100 mM TEAB, to shift pH for labeling and were tagged with stable-isotope dimethyl labels comprising regular formaldehyde and cyanoborohydride (28 Da shift, designated “light label”) or deuterated formaldehyde and regular cyanoborohydride (32 Da shift, designated “heavy label”) (all reagents from Sigma-Aldrich). Wild type samples were tagged with light labels and samples from *alg2^hypo^* mutant samples were tagged with heavy labels, including a label swap for one of the four eye sample replicates. Labeled peptides were washed with 20 μL of 0.1 % TFA and eluted from StageTip material by adding 10 μL of 50 % ACN in 0.1 % TFA twice with subsequent centrifugation for 1 min at 1500 g after each step. Differentially labeled samples were mixed in equal amounts, dried in a vacuum centrifuge and stored at −20 °C until LC-MS analysis.

#### LC-MS measurements

For technical reasons, whole hatchling and eye samples were analyzed using slightly different LC-MS setups. Approx. 5 and 8.3 μg of tryptic peptides per LC-MS injection were analyzed for hatchling samples (two technical replicate measurements per biological replicate with different amounts, respectively) and approx. 2 μg of tryptic peptides were analyzed for eye samples. A precursor ions inclusion list was used to improve run-to-run reproducibility for all samples. All used solvents were UPLC grade.

In brief, whole hatchling samples were resuspended after vacuum centrifuge in 20 % ACN / 0.1 % TFA and incubated for 5 min at RT, then diluted 10 fold with 0.1 % TFA prior to LC-MS measurement, which was conducted using an Ultimate 3000 UPLC (Thermo Fisher Scientific) coupled to a Q-Exactive HF mass spectrometer (Thermo Fisher Scientific). Analytical LC separation was performed using an in-house packed analytical column (20 cm length, 75 μm inner diameter; CS – Chromatographie Service GmbH) filled with 1.9 μm particle size, 120 Å pore size ReprosilPur-AQ 120 C18 material (Dr. Maisch) and carried out for 160 min total analysis time. The chromatographic method consisted of a linear gradient of buffer B (0.1 % *v*/*v* formic acid (FA), Proteochem, 10 % *v*/*v* H_2_O, Biosolve in ACN) in buffer A (0.1 % FA, 1 % ACN in H_2_O) from 3 to 40 % B in 120 min with a flow rate of 300 nL / min, followed by a washing (95 % B) and an equilibration step. Prior to the gradient, samples were loaded to the analytical column for 20 min with 3 % B at 550 nL / min flow rate. Eluting peptides were analyzed online by a coupled Q-Exactive-HF mass spectrometer running in DDA mode. Full scans were performed at 60 000 (m/z 200) resolution for a mass range covering 400-1600 m/z for 3e6 ions or up to a max IT of 45 ms. The full scan was followed by up to 15 MS/MS scans at 15 000 resolution with a max IT of 50 ms for up to 1e5 ions (AGC target). Precursors were isolated with a window of 1.6 m/z and fragmented with a collision energy of 27 (NCE). Unassigned and singly charged peptides were excluded from fragmentation and dynamic exclusion was set to 35 s.

For eye samples, dried peptides were resuspended in 2.5 % 1,1,1,3,3,3-Hexafluoro-2-propanol (Sigma-Aldrich) / 0.1 % TFA prior to LC-MS measurement, which was conducted using an Ultimate 3000 UPLC (Thermo Fisher Scientific) coupled to a Q-Exactive HF-X mass spectrometer (Thermo Fisher Scientific). During the LC separation, peptides were first loaded onto a trapping cartridge (Acclaim PepMap300 C18, 5 μm particle size, 300 Å pore size, Thermo Fisher Scientific) and washed for 3 min with 0.1 % TFA. Analytical separation was performed using a nanoEase MZ Peptide analytical column (BEH, 20 cm length, 75 μm inner diameter, 1.7 μm particle size, 300 Å pore size, Waters) and carried out for 150 min total analysis time. The chromatographic method consisted of a linear gradient of buffer B (0.1 % FA, 19.9 % H_2_O, Biosolve in ACN) in buffer A (0.1 % FA in H_2_O) from 5 to 38 % B in 132 min with a flow rate of 300 nL / min, followed by a washing (95 % B) and an equilibration step. Eluting peptides were analyzed online by a coupled Q-Exactive-HF-X mass spectrometer running in DDA mode. Full scans were performed at 60 000 resolution for a mass range covering 350-1500 m/z for 3e6 ions or up to a max IT of 45 ms. The full scan was followed by up to 20 MS/MS scans at 15 000 resolution with a max IT of 22 ms for up to 1e5 ions (AGC target). Precursors were isolated with a window of 1.6 m/z and fragmented with a collision energy of 27 (NCE). Unassigned and singly charged peptides were excluded from fragmentation and dynamic exclusion was set to 35 s.

#### Protein identification and relative quantification with MaxQuant

Raw files were processed using MaxQuant (version 1.6.12.0; (Cox & Mann, 2008)) for protein identification and quantification. MS / MS spectra were searched against the Uniprot Oryzias latipes database (retrieved in February 2020, last edited in November 2019), common contaminants and an additional fasta file containing the amino acid sequence of the usherin protein (Uniprot ID: U3R8H7) by Andromeda search engine with the following parameters: Carbamidomethylation of cysteine residues as fixed modification and Acetyl (Protein N-term), Oxidation (M) as variable modifications, trypsin/P as the proteolytic enzyme with up to two missed cleavages allowed. The maximum false discovery rate for proteins and peptides was 0.01 and a minimum peptide length of 7 amino acids was required. Match between runs and requantify options were disabled. Quantification mode was with the dimethyl Lys 0 and N-term 0 as light labels and dimethyl Lys 4 and N-term 4 as heavy labels. All other parameters were default parameters of MaxQuant. Quantitative normalized ratios were calculated by MaxQuant and used for further data analysis.

#### Statistical analysis of MS data

Perseus software (Version 1.5.6.0) was used (Tyanova *et al*, 2016). For the volcano plots in Fig 3, MaxQuant normalized (total protein normalization) and log2 transformed data were filtered (at least two razor or unique peptides per protein group required) and technical replicate values were averaged, if available. One-sample t-test (comparison to 0, p-value 0.05, −1< t-test difference < 1) was then performed and plots were produced with R and RStudio version 1.2.5042. For exclusive protein groups (only wt or only mutant), the raw intensity values for each biological replicate were normalized to the mean of unnormalized Gapdh ratios from all replicates and log2 transformed so that the proteins were plotted on the y axis according to their gapdh normalized intensities for representation only. Exclusive hits were only considered when present in at least two biological replicates.

## Acknowledgements

We thank Tanja Kellner for cloning and transcribing the sgRNAs used in this study. We thank Oi Pui Hoang and Alicia Perez for their initial contributions to start the project. We thank Antonino Saraceno, Erik Leist and Marcena Majewski for taking care of our fish. We thank Nicole Lübbehusen for excellent technical assistance and for performing dimethyl labeling. We thank the MS-based Protein Analysis Unit of the Genomics and Proteomics Core Facility at the German Cancer Research Center for assistance with sample measurement. We are grateful to Sabine Strahl and Patrick Neubert to share the original figure we modified for the discussion figure. We thank Udo Reichl for generously providing research infrastructure to ER.

Funded by the Deutsche Forschungsgemeinschaft (DFG, German Research Foundation) – Project-ID FOR2509 P04, P05, P09 and P10, – Forschungsgruppe FOR 2509.

## Author contributions

SG, TT and JW conceived the project which TT and JW supervising. TT designed the CRISPR/Cas9 strategy. SG performed all cloning, histology, staining, injections, mutant characterisation and other fish-related experiments and interpreted the results with TT and JW. VG performed and analyzed the xCGE-LIF-based *N*-glycomics. VG, MH and ER interpreted the *N*-glycomics data. RS and TR performed, curated and interpreted the proteomics data together with SG and TT. LB performed fibroblast experiments supervised by CT. TT, SG, RS, TR, VG and MH performed computational analysis. SG performed imaging and analysis. JW, TR, ER and CT provided resources. SG, TT and JW wrote the original draft with contributions of RS, TR, MH, VG, CT and ER. All authors reviewed and edited the manuscript finalized by SG, TT and JW.

## Data Availability Section

All relevant data are included in this manuscript and the extended view section.

## Expanded View Figures

**Figure EV1.**
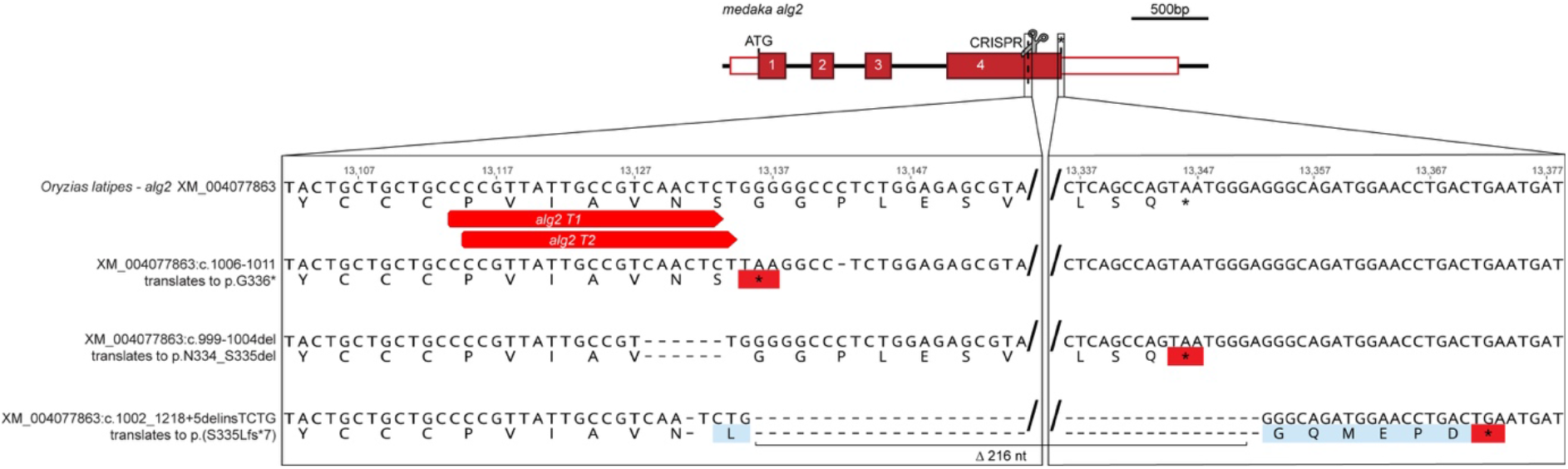
Genomic sequences of *alg2* alleles. Schematic representation of *Oryzias latipes alg2* locus with site of targeted mutagenesis indicated (grey scissors; red box, coding exons; white box, UTR). Zoom into the genomic sequence at the site of mutagenesis in wildtype and stable alleles of the three mutant *alg2* alleles. Potential aminoacid sequence given below codons of open reading frame. red, premature STOP, light blue, amino acids resulting from frameshift. Nomenclature according to (Dunnen *et al*, 2016).

**Figure EV2.**
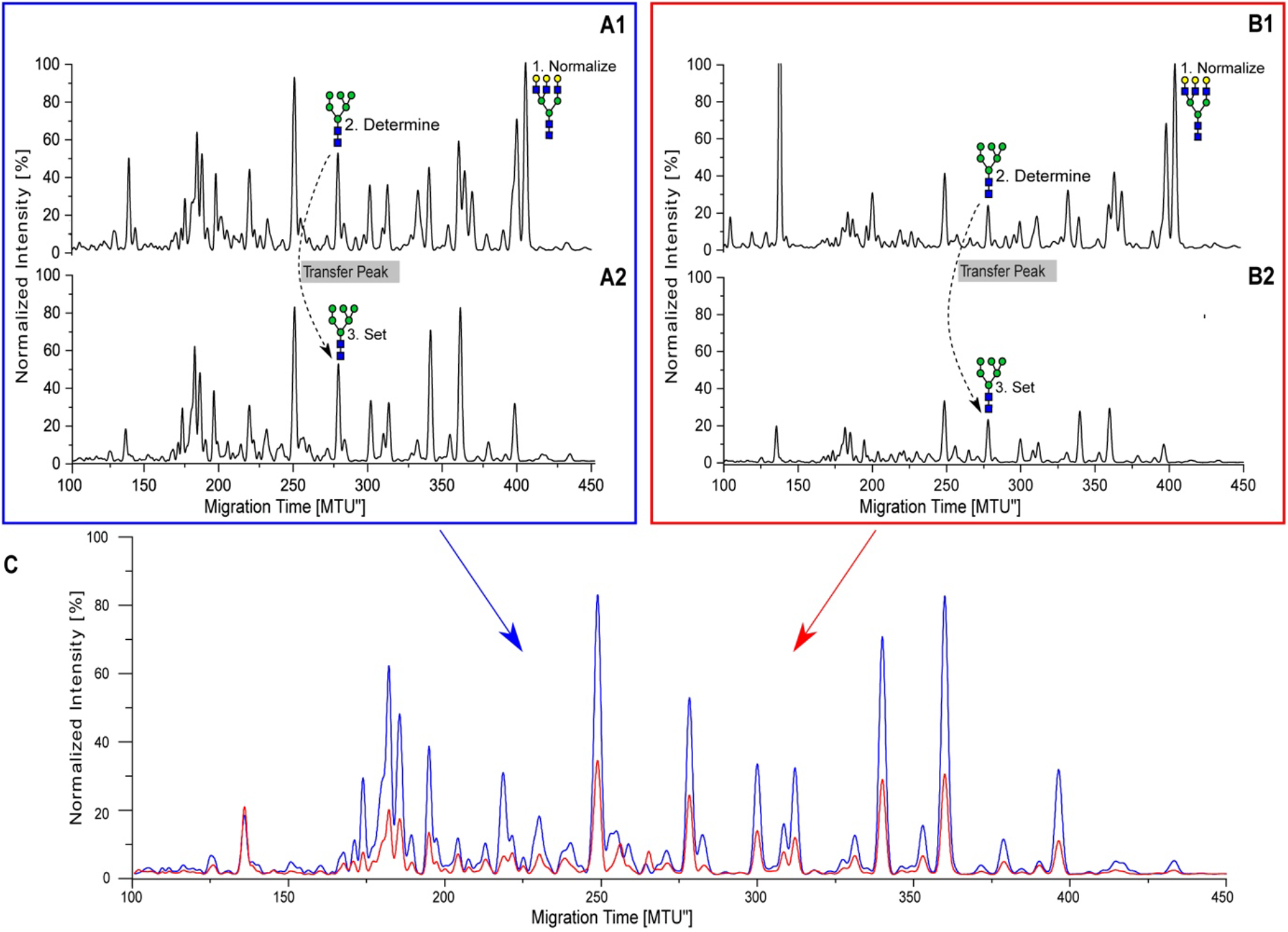
Normalization of xCGE-LIF electropherogram. Schematic overview of the quantitative normalization for *N*-glycan fingerprints using an internal standard. A1/B1: *N*-glycan fingerprints of sample A and B spiked with internal standard (bovine asialofetuin). Intensities are normalized to the asialofetuin-derived *N*-glycan peak (A3G3). A2/B2: A sample-derived *N*-glycan peak (Man6) is used to transfer the quantitative normalization to *N*-glycan fingerprints not containing the internal standard. C: Overlay of two samples (A2 and B2) comprising a quantitative normalization enabling inter-sample comparison.

**Figure EV3.**
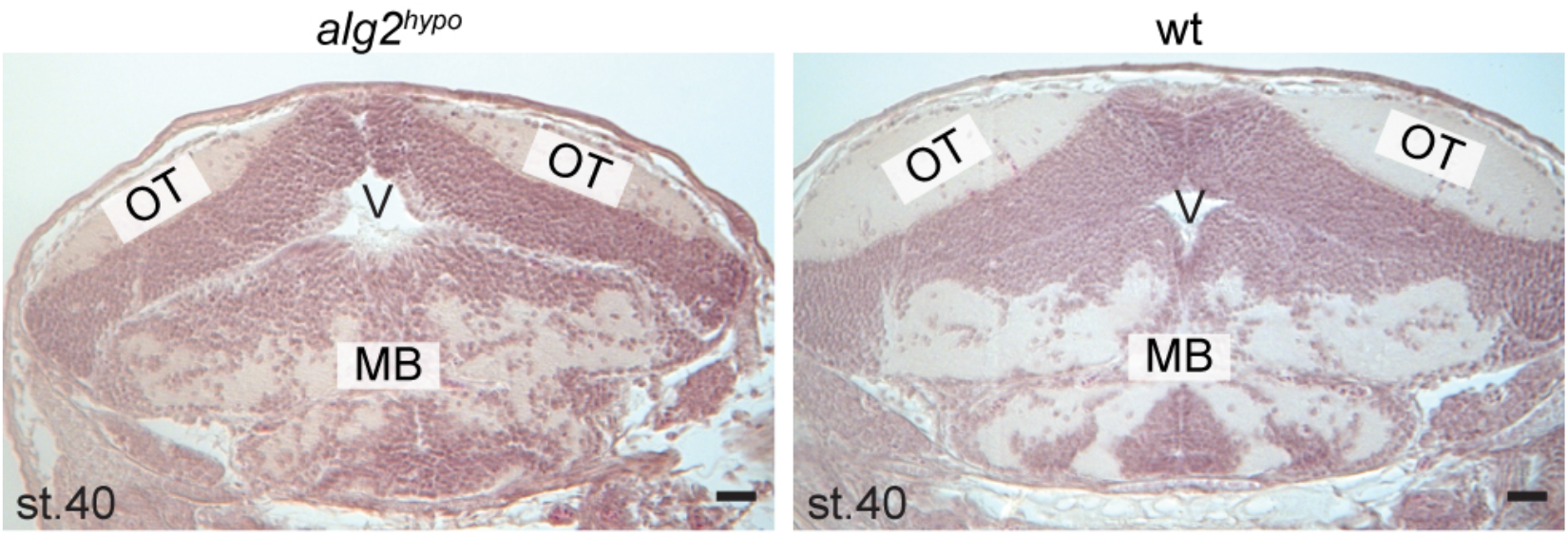
*Alg2^hypo^* homozygotes show reduced white matter. Hematoxylin and eosin staining on transverse sections of *alg2^hypo^* (left) and wild type (wt) embryos (right) at stage 40. Note overall reduction of white matter most pronounced in the optic tectum (OT). Larger ventricle (V) and not well confined lining as compared to wt. Scale bar: 50 μm

**Expanded View Table 1**

*N*-glycan structures identified in lysates from medaka wild type and *alg2^hypo^* hatchlings. Relative intensities are based on normalization to internal standard. Symbolic representation of *N*-glycan structures were drawn with GlycoWorkbench Version 1.1, following the guideline of the Consortium for Functional Glycomics (Varki *et al*, 2009). Glycan names are adapted from the Oxford nomenclature.

**Expanded View Table 2**

Proteomics results of stage 40 whole hatchlings differential proteomics *alg2^hypo^*/wt (sheet 1) and exclusive hits (sheet 2) and enucleated eye samples differential proteomics *alg2^hypo^*/wt (sheet 3) and exclusive hits (sheet 4).

## References

Ba-Abbad R, Leys M, Wang X, Chakarova C, Waseem N, Carss KJ, Raymond FL, Bujakowska KM, Pierce EA, Mahroo OA, Mohamed MD, Holder GE, Hummel M, Arno G & Webster AR (2018) Clinical Features of a Retinopathy Associated With a Dominant Allele of the RGR Gene. Invest. Ophthalmol. Vis. Sci. 59:4812–4820

Boersema PJ, Raijmakers R, Lemeer S, Mohammed S & Heck AJR (2009) Multiplex peptide stable isotope dimethyl labeling for quantitative proteomics. Nature Protocols 4:484–494

Breitling J & Aebi M (2013) N-linked protein glycosylation in the endoplasmic reticulum. Cold Spring Harbor Perspectives in Biology 5:a013359–a013359

Burns DM & Touster O (1982) Purification and characterization of glucosidase II, an endoplasmic reticulum hydrolase involved in glycoprotein biosynthesis. J. Biol. Chem. 257:9990–10000

Cantagrel V, Lefeber DJ, Ng BG, Guan Z, Silhavy JL, Bielas SL, Lehle L, Hombauer H, Adamowicz M, Swiezewska E, De Brouwer AP, Blümel P, Sykut-Cegielska J, Houliston S, Swistun D, Ali BR, Dobyns WB, Babovic-Vuksanovic D, van Bokhoven H, Wevers RA, et al (2010) SRD5A3 is required for converting polyprenol to dolichol and is mutated in a congenital glycosylation disorder. Cell 142:203–217

Carrigan M, Duignan E, Humphries P, Palfi A, Kenna PF & Farrar GJ (2016) A novel homozygous truncating GNAT1 mutation implicated in retinal degeneration. Br J Ophthalmol 100:495–500

Centanin L, Ander J-J, Hoeckendorf B, Lust K, Kellner T, Kraemer I, Urbany C, Hasel E, Harris WA, Simons BD & Wittbrodt J (2014) Exclusive multipotency and preferential asymmetric divisions in post-embryonic neural stem cells of the fish retina. Development 141:3472–3482

Ceroni A, Maass K, Geyer H, Geyer R, Dell A & Haslam SM (2008) GlycoWorkbench: a tool for the computer-assisted annotation of mass spectra of glycans. J. Proteome Res. 7:1650–1659

Chang IJ, He M & Lam CT (2018) Congenital disorders of glycosylation. Ann Transl Med 6: 477–477

Cline A, Gao N, Flanagan-Steet H, Sharma V, Rosa S, Sonon R, Azadi P, Sadler KC, Freeze HH, Lehrman MA & Steet R (2012) A zebrafish model of PMM2-CDG reveals altered neurogenesis and a substrate-accumulation mechanism for N-linked glycosylation deficiency. Mol. Biol. Cell 23:4175–4187

Conley SM, Stuck MW, Watson JN & Naash MI (2017) Rom1 converts Y141C-Prph2-associated pattern dystrophy to retinitis pigmentosa. Human Molecular Genetics 26:509–518

Cossins J, Belaya K, Hicks D, Salih MA, Finlayson S, Carboni N, Liu WW, Maxwell S, Zoltowska K, Farsani GT, Laval S, Seidhamed MZ, WGS500 Consortium, Donnelly P, Bentley D, McGowan SJ, Müller J, Palace J, Lochmüller H & Beeson D (2013) Congenital myasthenic syndromes due to mutations in ALG2 and ALG14. Brain 136:944–956

Cox J & Mann M (2008) MaxQuant enables high peptide identification rates, individualized p.p.b.-range mass accuracies and proteome-wide protein quantification. Nat. Biotechnol. 26:1367–1372

Daenzer JMI, Sanders RD, Hang D & Fridovich-Keil JL (2012) UDP-galactose 4’-epimerase activities toward UDP-Gal and UDP-GalNAc play different roles in the development of Drosophila melanogaster. PLoS Genet 8:e1002721

Dunnen den JT, Dalgleish R, Maglott DR, Hart RK, Greenblatt MS, McGowan-Jordan J, Roux A-F, Smith T, Antonarakis SE & Taschner PEM (2016) HGVS Recommendations for the Description of Sequence Variants: 2016 Update. Hum. Mutat. 37:564–569

Ferrari S, Di Iorio E, Barbaro V, Ponzin D, Sorrentino FS & Parmeggiani F (2011) Retinitis pigmentosa: genes and disease mechanisms. Curr. Genomics 12:238–249

Gao X-D, Nishikawa A & Dean N (2004) Physical interactions between the Alg1, Alg2, and Alg11 mannosyltransferases of the endoplasmic reticulum. Glycobiology 14:559–570

Grinna LS & Robbins PW (1979) Glycoprotein biosynthesis. Rat liver microsomal glucosidases which process oligosaccharides. J. Biol. Chem. 254:8814–8818

Grinna LS & Robbins PW (1980) Substrate specificities of rat liver microsomal glucosidases which process glycoproteins. J. Biol. Chem. 255:2255–2258

Gupta R, Jung E, Work SBRTU2016 Prediction of N-glycosylation sites in human proteins. 2004

Gutierrez-Triana JA, Tavhelidse T, Thumberger T, Thomas I, Wittbrodt B, Kellner T, Anlas K, Tsingos E & Wittbrodt J (2018) Efficient single-copy HDR by 5 modified long dsDNA donors. eLife 7:142

Haeuptle MA & Hennet T (2009) Congenital disorders of glycosylation: an update on defects affecting the biosynthesis of dolichol-linked oligosaccharides. Hum. Mutat. 30:1628–1641

Helenius A & Aebi M (2004) Roles of N-linked glycans in the endoplasmic reticulum. Annu. Rev. Biochem. 73:1019–1049

Hennet T & Cabalzar J (2015) Congenital disorders of glycosylation: a concise chart of glycocalyx dysfunction. Trends Biochem. Sci. 40:377–384

Hennig R, Cajic S, Borowiak M, Hoffmann M, Kottler R, Reichl U & Rapp E (2016) Towards personalized diagnostics via longitudinal study of the human plasma N-glycome. Biochim. Biophys. Acta 1860:1728–1738

Iwamatsu T (2004) Stages of normal development in the medaka Oryzias latipes. Mech. Dev. 121:605–618

Kaushal S, Ridge KD & Khorana HG (1994) Structure and function in rhodopsin: the role of asparagine-linked glycosylation. PNAS 91:4024–4028

Khoriaty R, Vasievich MP & Ginsburg D (2012) The COPII pathway and hematologic disease. Blood 120:31–38

Lee AS (2005) The ER chaperone and signaling regulator GRP78/BiP as a monitor of endoplasmic reticulum stress. METHODS 35:373–381

Li S-T, Wang N, Xu X-X, Fujita M, Nakanishi H, Kitajima T, Dean N & Gao X-D (2018) Alternative routes for synthesis of N-linked glycans by Alg2 mannosyltransferase. FASEB J. 32:2492–2506

Lust K & Wittbrodt J (2018) Activating the regenerative potential of Müller glia cells in a regeneration-deficient retina. eLife 7:7028

Matthijs G, Schollen E, Van Schaftingen E, Cassiman JJ & Jaeken J (1998) Lack of homozygotes for the most frequent disease allele in carbohydrate-deficient glycoprotein syndrome type 1A. Am. J. Hum. Genet. 62:542–550

Med HFCCL2002 Congenital disorders of glycosylation: an emerging group of inherited diseases with multisystemic clinical presentation

Meunier L, Usherwood Y-K, Chung KT & Hendershot LM (2002) A subset of chaperones and folding enzymes form multiprotein complexes in endoplasmic reticulum to bind nascent proteins. Mol. Biol. Cell 13:4456–4469

Monies D, Alhindi HN, Almuhaizea MA, Abouelhoda M, Alazami AM, Goljan E, Alyounes B, Jaroudi D, AlIssa A, Alabdulrahman K, Subhani S, El-Kalioby M, Faquih T, Wakil SM, Altassan NA, Meyer BF & Bohlega S (2016) A first-line diagnostic assay for limb-girdle muscular dystrophy and other myopathies. Hum. Genomics 10:32–7

Morava E, Wevers RA, Cantagrel V, Hoefsloot LH, Al-Gazali L, Schoots J, van Rooij A, Huijben K, van Ravenswaaij-Arts CMA, Jongmans MCJ, Sykut-Cegielska J, Hoffmann GF, Bluemel P, Adamowicz M, van Reeuwijk J, Ng BG, Bergman JEH, van Bokhoven H, Körner C, Babovic-Vuksanovic D, et al (2010) A novel cerebello-ocular syndrome with abnormal glycosylation due to abnormalities in dolichol metabolism. Brain 133:3210–3220

Muntoni F, Torelli S & Brockington M (2008) Muscular dystrophies due to glycosylation defects. Neurotherapeutics 5:627–632

Murray AR, Fliesler SJ & Al-Ubaidi MR (2009) Rhodopsin: the functional significance of asn-linked glycosylation and other post-translational modifications. Ophthalmic Genet. 30:109–120

Oki T, Yamazaki K, Kuromitsu J, Okada M & Tanaka I (1999) cDNA cloning and mapping of a novel subtype of glutamine:fructose-6-phosphate amidotransferase (GFAT2) in human and mouse. Genomics 57:227–234

Park C & Zhang J (2011) Genome-wide evolutionary conservation of N-glycosylation sites. Mol. Biol. Evol. 28:2351–2357

Rappsilber J & Mann M (2007) Analysis of the topology of protein complexes using cross-linking and mass spectrometry. CSH Protoc 2007:pdb.prot4594–pdb.prot4594

Reinhardt R, Centanin L, Tavhelidse T, Inoue D, Wittbrodt B, Concordet J-P, Martinez-Morales JR & Wittbrodt J (2015) Sox2, Tlx, Gli3, and Her9 converge on Rx2 to define retinal stem cells in vivo. The EMBO Journal 34:1572–1588

Rupp RA, Snider L & Weintraub H (1994) Xenopus embryos regulate the nuclear localization of XMyoD. Genes Dev. 8:1311–1323

Schindelin J, Arganda-Carreras I, Frise E, Kaynig V, Longair M, Pietzsch T, Preibisch S, Rueden C, Saalfeld S, Schmid B, Tinevez J-Y, White DJ, Hartenstein V, Eliceiri K, Tomancák P & Cardona A (2012) Fiji: an open-source platform for biological-image analysis. Nat Meth 9:676–682

Sharma V, Nayak J, DeRossi C, Charbono A, Ichikawa M, Ng BG, Grajales-Esquivel E, Srivastava A, Wang L, He P, Scott DA, Russell J, Contreras E, Guess CM, Krajewski S, Del Rio-Tsonis K & Freeze HH (2014) Mannose supplements induce embryonic lethality and blindness in phosphomannose isomerase hypomorphic mice. FASEB J. 28:1854–1869

Stemmer M, Thumberger T, del Sol Keyer M, Wittbrodt J & Mateo JL (2015) CCTop: An Intuitive, Flexible and Reliable CRISPR/Cas9 Target Prediction Tool. PLoS ONE 10:e0124633

Sullivan LS, Bowne SJ, Koboldt DC, Cadena EL, Heckenlively JR, Branham KE, Wheaton DH, Jones KD, Ruiz RS, Pennesi ME, Yang P, Davis-Boozer D, Northrup H, Gurevich VV, Chen R, Xu M, Li Y, Birch DG & Daiger SP (2017) A Novel Dominant Mutation in SAG, the Arrestin-1 Gene, Is a Common Cause of Retinitis Pigmentosa in Hispanic Families in the Southwestern United States. Invest. Ophthalmol. Vis. Sci. 58:2774–2784

Szegezdi E, Logue SE, Gorman AM & Samali A (2006) Mediators of endoplasmic reticulum stress-induced apoptosis. EMBO Rep. 7:880–885

Szklarczyk D, Gable AL, Lyon D, Junge A, Wyder S, Huerta-Cepas J, Simonovic M, Doncheva NT, Morris JH, Bork P, Jensen LJ & Mering CV (2019) STRING v11: protein-protein association networks with increased coverage, supporting functional discovery in genome-wide experimental datasets. Nucleic Acids Research 47:D607–D613

Tam BM & Moritz OL (2009) The role of rhodopsin glycosylation in protein folding, trafficking, and light-sensitive retinal degeneration. Journal of Neuroscience 29:15145–15154

Thiel C, Lübke T, Matthijs G, Figura von K & Körner C (2006) Targeted disruption of the mouse phosphomannomutase 2 gene causes early embryonic lethality. Molecular and Cellular Biology 26:5615–5620

Thiel C, Schwarz M, Peng J, Grzmil M, Hasilik M, Braulke T, Kohlschütter A, Figura von K, Lehle L & Körner C (2003) A new type of congenital disorders of glycosylation (CDG-Ii) provides new insights into the early steps of dolichol-linked oligosaccharide biosynthesis. J. Biol. Chem. 278:22498–22505

Thiesler CT, Cajic S, Hoffmann D, Thiel C, van Diepen L, Hennig R, Sgodda M, Weiβmann R, Reichl U, Steinemann D, Diekmann U, Huber NMB, Oberbeck A, Cantz T, Kuss AW, Körner C, Schambach A, Rapp E & Buettner FFR (2016) Glycomic Characterization of Induced Pluripotent Stem Cells Derived from a Patient Suffering from Phosphomannomutase 2 Congenital Disorder of Glycosylation (PMM2-CDG). Mol. Cell Proteomics 15:1435–1452

Tyanova S, Temu T, Sinitcyn P, Carlson A, Hein MY, Geiger T, Mann M & Cox J (2016) The Perseus computational platform for comprehensive analysis of (prote)omics data. Nat Meth 13:731–740

Varki A, Cummings R, Esko J, Freeze H & Search GHI-19 (2019) Essentials of Glycobiology. 2nd ed2009

Varki A, Cummings RD, Esko JD, Freeze HH, Stanley P, Marth JD, Bertozzi CR, Hart GW & Etzler ME (2009) Symbol nomenclature for glycan representation. Proteomics 9:5398–5399

Wessel D & Flügge UI (1984) A method for the quantitative recovery of protein in dilute solution in the presence of detergents and lipids. Anal. Biochem. 138:141–143

Zecha J, Meng C, Zolg DP, Samaras P, Wilhelm M & Kuster B (2018) Peptide Level Turnover Measurements Enable the Study of Proteoform Dynamics. Mol. Cell Proteomics 17:974–992

Zuber C, Spiro MJ, Guhl B, Spiro RG & Roth J (2000) Golgi apparatus immunolocalization of endomannosidase suggests post-endoplasmic reticulum glucose trimming: implications for quality control. Mol. Biol. Cell 11:4227–4240

